# Dissecting Branch-Specific Unfolded Protein Response Activation in Drug-Tolerant BRAF-Mutant Melanoma using Data-Independent Acquisition Mass Spectrometry

**DOI:** 10.1101/2025.03.20.644425

**Authors:** Lea A. Barny, Jake N. Hermanson, Sarah K. Garcia, Philip E. Stauffer, Lars Plate

## Abstract

Cells rely on the Unfolded Protein Response (UPR) to maintain ER protein homeostasis (proteostasis) when faced with elevated levels of misfolded and aggregated proteins. The UPR is comprised of three main branches—ATF6, IRE1, and PERK—that coordinate the synthesis of proteins involved in folding, trafficking, and degradation of nascent proteins to restore ER function. Dysregulation of the UPR is linked to numerous diseases, including neurodegenerative disorders, cancer, and diabetes. Despite its importance, identifying UPR targets has been challenging due to their heterogeneous induction, which varies by cell type and tissue. Additionally, defining the magnitude and range of UPR-regulated genes is difficult because of intricate temporal regulation, feedback between UPR branches, and extensive cross-talk with other stress-signaling pathways. To comprehensively identify UPR-regulated proteins and determine their branch specificity, we developed a data-independent acquisition (DIA) liquid-chromatography mass spectrometry (LC-MS) pipeline. Our optimized workflow improved identifications of low-abundant UPR proteins and leveraged an automated SP3-based protocol on the Biomek i5 liquid handler for label-free peptide preparation. Using engineered stable cell lines that enable selective pharmacological activation of each UPR branch without triggering global UPR activation, we identified branch-specific UPR proteomic targets. These targets were subsequently applied to investigate proteomic changes in multiple patient-derived BRAF-mutant melanoma cell lines treated with a BRAF inhibitor (PLX4720, i.e., vemurafenib). Our findings revealed differential regulation of the XBP1s branch of the UPR in the BRAF-mutant melanoma cell lines after PLX4720 treatment, likely due to calcium activation, suggesting that the UPR plays a role as a non-genetic mechanism of drug tolerance in melanoma. In conclusion, the validated branch-specific UPR proteomic targets identified in this study provide a robust framework for investigating this pathway across different cell types, drug treatments, and disease conditions in a high-throughput manner.

## Introduction

The endoplasmic reticulum (ER) plays a critical role in the processing of secretory proteins, which comprise about one-third of the human proteome^1^. When ER protein folding capacity is compromised or ER luminal conditions become unfavorable (altered redox state, dysregulated calcium levels, increased reactive oxygen species, etc.), “ER stress” results from an accumulation of aggregated and/or misfolded proteins in the ER. The Unfolded Protein Response (UPR) mitigates ER stress through a highly conserved and adaptive signal transduction pathway^2–4^ mediated by three integrated signaling cascades named after their respective ER transmembrane stress-sensing proteins: ATF6 (Activating Transcription Factor 6), IRE1 (Inositol Requiring Enzyme 1), and PERK (Protein Kinase RNA-like ER kinase) (**Fig. 1A**). Activation of ATF6, IRE1, and PERK leads to the downstream production of their corresponding transcription factors (TFs): p50ATF6, XBP1s, and ATF4, respectively. Each UPR associated TF is generated through a distinct mechanism: regulated proteolysis, nonconventional mRNA splicing, and translational control, respectively^2,5^. Ultimately, binding of UPR-associated TFs to DNA results in the expression of a network of genes to restore ER protein homeostasis (proteostasis) or promote induction of apoptosis when ER stress is unresolved^6^.

**Figure 1.**
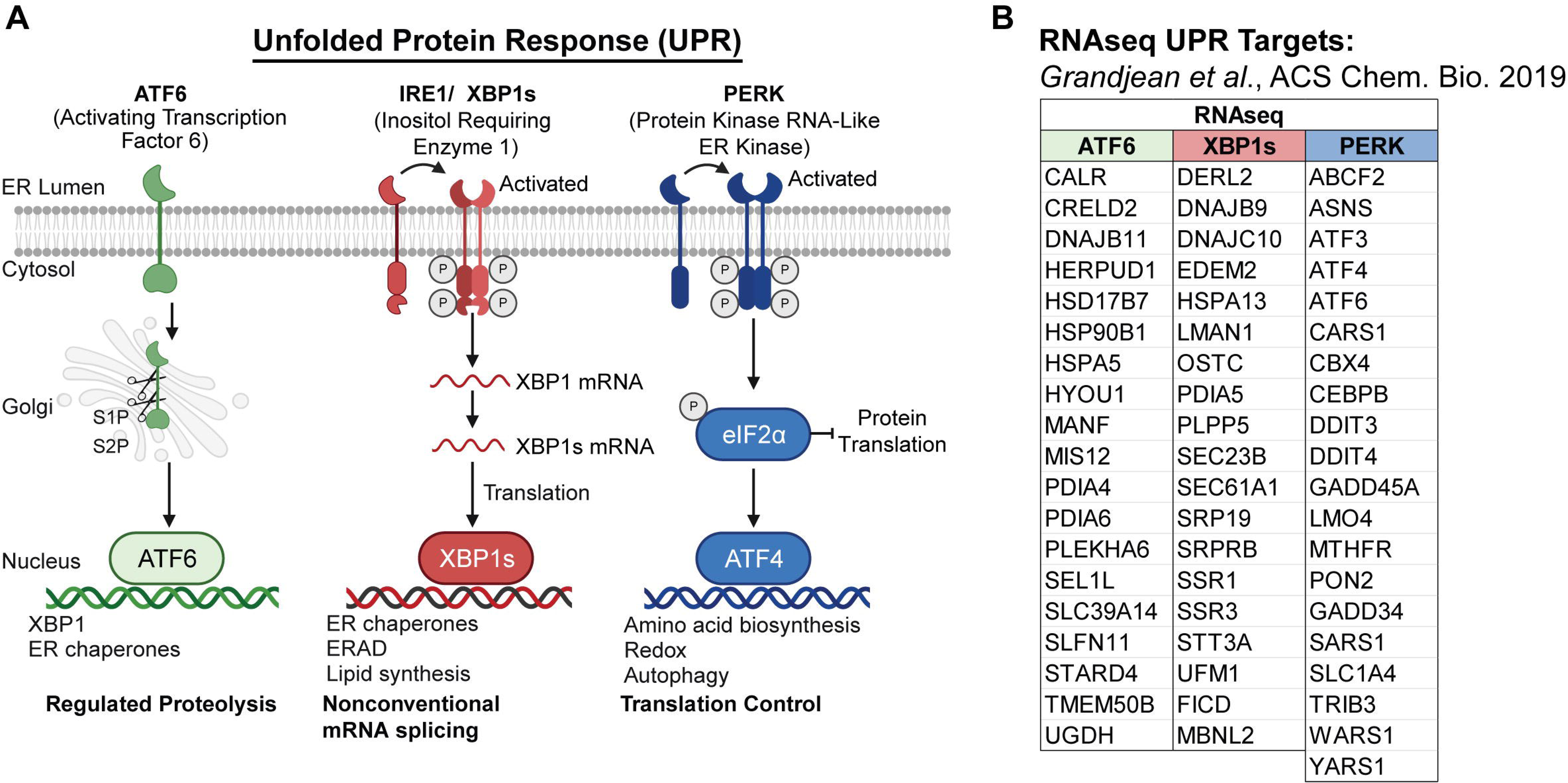
UPR overview. (**A**) Schematic showing the three major branches of the UPR (ATF6, XBP1s, and PERK) along with their respective transcription factors and the previously described transcriptional and translational remodeling induced by each arm of the response. (**B**) Previously elucidated transcriptional targets of each branch of the UPR.^6^

The integrity of secretory proteostasis is maintained by balancing two major ER quality control processes: protein folding and degradation^7^. An increase in ER folding capacity is primarily achieved through the transcriptional and translational remodeling of ER proteostasis pathways by ATF6 and IRE1. Transcriptional targets of XBP1s include those involved in all aspects of the UPR: protein folding (i.e. ATP-dependent chaperones, lectin chaperones, protein disulfide isomerases, and other folding factors), degradation (ER-associated degradation or ERAD), ER protein import, N-linked glycosylation, and vesicular trafficking^8,9^. On the other hand, p50ATF6 mainly induces the expression of protein folding machinary^10^. XBP1s and p50ATF6 can also either homo- or heterodimerize to induce distinct genes involved in overlapping ER quality control pathways^10,11^. Moreover, the UPR modulates ER protein folding load by translational modulation. PERK activation, which results in the phosphorylation of eukaryotic initiation factor 2 alpha subunit (eIF2α), leads to transient translation attenuation to decrease the influx of proteins entering the ER for folding^4,12^. Additional kinases can also mediate phosphorylation of eIF2α, in a pathway known as the integrated stress response (ISR)^13^. Regardless of which kinase phosphorylates eIF2α, transcription factors ATF4 and CHOP act as the main regulators of the PERK branch of the UPR and the ISR^14^. This results in increased expression of oxidative stress genes and ultimately pro-apoptotic genes, if the ER stress is not resolved^15–17^ . Typically, over the course of the response, negative feedback from GADD34 results in the dephosphorylation of eIF2α reinitiating translation and restoring normal protein synthesis.

Dysregulation of the UPR plays a role in numerous pathologies, including neurodegenerative diseases, cancer, and diabetes^18^. For instance, chronic ER stress in pancreatic beta-cells leads to cell death and the onset of diabetes^19–21^. Furthermore, a diminished capacity to activate the UPR during aging is associated with a reduced capacity to clear misfolded and aggregated protein, thereby accelerating the onset of neurodegenerative diseases^22,23^. Many cancers also rely on prolonged UPR activation to maintain high protein folding and cell proliferation rates^24,25^. Thus, understanding the regulation of the UPR in the context of disease states is critical to elucidating disease mechanisms and developing pharmaceutical interventions.

Approximately half of all melanoma patients harbor a pro-oncogenic BRAF mutation^26^. However, BRAF mutations are also common in various other cancer types^27^. When BRAF carries the missense mutation V600E (or V600K/ V600R) within its kinase loop, it leads to constitutive activation of the kinase and subsequently the MAPK pathway. This dysregulation results in altered cell cycle progression, increased cell proliferation, altered metabolism, enhanced migration, disrupted differentiation, and evasion of apoptosis^28,29^. BRAF-mutant melanoma is commonly treated with BRAF inhibitors (BRAFi)^30,31^, however drug tolerance and eventual resistance is common. Upon initial drug exposure, BRAFi treated melanoma cells phenotypically adapt and exhibit a non-quiescent, slowly dividing state which is marked by balanced cell death and division (zero net growth), previously described as “idling”. This phenotype is derived from non-genetic mechanisms of drug response, as it can be reversed upon discontinuation of BRAFi treatment^32,33^. Activated calcium signaling in these idling cells is thought to serve as a major mechanism of non-genetic adaptation resulting in reactivation of the MAPK pathway, which may coordinate with the activation of the UPR to further promote drug-tolerance^34^. Hence, we sought to characterize UPR activation in this context.

The induction of UPR targets are heterogeneous and can vary by cell type and/or tissue^35,36^. Defining both the induction magnitude and the repertoire of UPR-regulated genes remains challenging because of intricate temporal regulation, feedback between UPR pathways, and a high degree of cross-talk between the UPR and other stress-signaling pathways, such as the oxidative stress-response (OSR), the integrated stress-response pathway (ISR), or the mitochondrial unfolded protein response (UPR^mito^) pathways^37,38^. The intricacies and heterogeneity in UPR regulation at the proteomics level often remain hidden as the UPR is typically characterized by quantifying only select protein markers (by Western blot) or by transcriptomics analysis, which lacks information on translational regulation.

Previous work by Grandjean and colleagues defined branch-specific UPR transcriptional targets (**Fig. 1B; Supplemental Table S1**)^6^. Integrating these gene sets into a targeted RNAseq analysis enabled the accurate deconvolution of the stress pathways activated by pharmacological agents. Here, we sought to develop a complementary proteomics platform to quantify UPR activation and branch-specific regulation. We took advantage of the detection sensitivity of data-independent acquisition (DIA) mass spectrometry since many of the proteins regulated by the UPR are lowly abundant and have only been detected stochastically in previous data-dependent acquisition studies. For example, prior investigations into the pharmacological manipulation of the pathway^39,40^ and how it differs between disease states^23,41,42^ have relied heavily on TMT-DDA methodologies or stable isotope labeling by amino acids in cell culture (SILAC)-DDA. Our DIA LC-MS/MS approach allowed us to refine and expand the list of proteostasis factors previously described in RNAseq datasets as preferentially regulated by the three distinct branches of the UPR.

To assess the utility of our branch-specific proteomics targets in a disease model, we applied our methodology to study the UPR in BRAF-mutant melanoma cell lines. When these cell lines were challenged with BRAF inhibitors, our analysis uncovered differential use of the UPR, with the IRE1/ XBP1s branch being especially heterogenous in its degree of activation. Overall, our work provides a valuable resource for evaluating branch-specific UPR activation via LC-MS/MS proteomics to facilitate comprehensive characterization of the UPR in different cell-types and disease contexts.

## Experimental Procedures

### Experimental Design and Statistical Rationale

Four biological replicates per drug treatment of pharmacologically activated HEK293^DAX^ and Fv2e-PERK were measured with DIA LC-MS/MS analysis. For MS acquisition comparison, identical pharmacologically activated HEK293^DAX^ and Fv2e-PERK cell lysate as was measured by DIA-MS was prepared as a single 18-plex for TMT-DDA analysis containing three biological replicates per each drug treatment (3x DMSO HEK293^DAX^, 3x TMP HEK293^DAX^, 3x DOX HEK293^DAX^, 3x DMSO Fv2e-PERK, 3x AP20187 (6 h) Fv2e-PERK, and 3x AP20187 (18 h) Fv2e-PERK). For validation, four biological replicates of global UPR activated samples (DMSO/ Tm/ Tg) were measured via DIA LC-MS/MS analysis. Lastly, four BRAF-mutant melanoma cell lines (A375, SKMEL5, SKMEL28, and WM88) treated with either vehicle (0.1% DMSO) or PLX4720 (10 µM) were collected after 3 or 8 days of drug treatment then prepared for MS analysis. In total, 64 BRAF-mutated melanoma samples were measured via DIA LC-MS/MS. When samples were prepared on the Biomek i5, 20 µg of *E. coli* lysate was digested and analyzed via DIA LC-MS/MS to ensure the quality of the digestion. Additionally, prior to the start of each MS analysis, 600 ng of *E. coli* peptides were measured with unfractionated DIA to ensure consistency of instrument performance. Retention time standards were not utilized. Statistical tests used to process proteomics data can be found in the experimental procedures section titled: “Data Analysis (Proteomics)”.

### Cell Lines with respective drug treatments

#### HEK293^DAX^, HEK293 Fv2e-PERK, and HEK293

Two engineered stable cell lines (HEK293^DAX^ and HEK293 Fv2e-PERK) which exhibit branch-specific transcriptional and translational reprogramming of the UPR in the absence of global ER stress, were provided by the Wiseman laboratory (The Scripps Research Institute, La Jolla, CA, USA). For experiments, cells were seeded at 1.2 million cells total in 6 cm plates. HEK293^DAX^ cells were treated with either dimethyl sulfoxide (0.1% v/v; DMSO; Sigma), trimethoprim (TMP; 10 µM; Fischer), doxycycline (DOX; 1 µg/mL; Sigma), or both, TMP (10 µM) and DOX (1 µg/ml), for 16 hours. TMP and DOX treatment upregulate the ATF6 and XBP1s transcription factors, respectively. Fv2e-PERK cells were treated with AP2017 (5 nM; Sigma) for either 6 or 18 h and DMSO (0.1% v/v) for 18 h to induce dimerization of PERK, resulting in eIF2α phosphorylation. Alternatively, to induce global UPR activation, Thapsigargin (Tg; 500 nM; Cayman Chemical) and Tunicamycin (Tm; 500 nM; Cayman Chemical) were applied to HEK293 cells for 16 h. All cells were cultured in Dulbecco’s modified Eagle medium (DMEM) supplemented with 10% fetal bovine serum (FBS), 1% penicillin/streptomycin (P/S), and 1% glutamine (Q). Cells were incubated at 37°C at 5% CO_2_. Experiments were performed on cells negative for mycoplasma with passage numbers between 20-30.

#### BRAF-Mutant Melanoma

Four BRAF-mutant melanoma cell lines (A375, SKMEL5, SKMEL28, and WM88) were obtained from the Vito Quaranta laboratory at Vanderbilt University and cultured in DMEM / F12 (Gibco) supplemented with 10% FBS and 1% P/S. Cells were maintained at 37°C in a 5% CO2 incubator, following previously described methods^34^. For drug treatment experiments, melanoma cells were detached from 10 cm culture dishes using TrypLE (Thermo Fisher) and seeded into 6-well plates. Twelve hours after seeding, cells were treated with PLX4720 (MedChemExpress), a research analog of vemurafenib^43,44^. A375, SKMEL5, and SKMEL28 cell lines were treated with 8 µM PLX4720, while the WM88 cell line was treated with 1 µM PLX4720. Each cell line was additionally drug-treated with vehicle, 0.1% DMSO, for 3 and 8 days as a control. Fresh vehicle and PLX4720 treatment media were applied to cells every 3 days until collection. Cell seeding densities were optimized to achieve 80-90% confluency upon collection at both the 3 and 8 day timepoint, with and without drug treatment. A total of four biological replicates were prepared for each cell line, with and without drug treatment at both time points, resulting in 64 total samples.

### Protein Collection

#### HEK293^DAX^, HEK293 Fv2e-PERK, and HEK293

Cells were collected via cell scraping. Briefly, cells were washed twice with cold phosphate buffered saline (PBS): 50 mM Tris (pH 7.5), 150 mM NaCl, 0.1% SDS, 1% Triton X-100, 0.5% deoxycholate) then scraped in PBS supplemented with 1 mM ethylenediaminetetraacetic acid (ETDA), followed by cell pelleting (200 g, 5 min). The pellet was then washed twice with cold PBS. Cell pellets were lysed in radioimmunoprecipitation assay (RIPA) buffer (50 mM Tris (pH 7.5), 150 mM NaCl, 0.1% SDS, 1% Triton X-100, 0.5% deoxycholate) with Roche complete EDTA-free protease inhibitor (Sigma) for 30 min total on ice (vortexing at 15 min). Fv2e-PERK cells were additionally lysed with 1% (v/v) phosphate inhibitor (Sigma). Cells were subsequently centrifuged for 15 min at 21.1 kg to remove cellular debris. The lysate was then transferred to fresh tubes and the protein concentration was quantified using a bicinchoninic acid assay (BCA; ThermoFisher Scientific)^45^.

#### BRAF-Mutant Melanoma

Cells were placed on ice and washed twice with cold PBS. After washing, cells were lysed directly on the tissue culture plate using RIPA buffer at 4°C (rocking) for 30 minutes. Following lysis, the cells were scraped into 1.5 ml centrifuge tubes and centrifuged at 16,000 g for 15 minutes. The cleared lysate was then used for a BCA assay to determine the protein concentration.

#### Western Blot Analysis (Drug Treated HEK293^DAX^, HEK293 Fv2e-PERK, HEK293)

To validate samples prior to LC-MS/MS analysis, 6x Laemmli buffer (12% SDS, 125 mM Tris, pH 6.8, 20% glycerol, bromophenol blue, 100 mM DDT) was added to 20 µg of protein per sample and heated for 5 min at 95°C. Samples were then separated on an 12% SDS-PAGE gel (60 V for 20 min, 160 V for 80 min) and transferred to a 0.45 µm PVDF membrane (100 V for 80 min;Sigma) for Western blotting. Prior to blotting, the blot was blocked with 5% (w/v) molecular-grade non-fat milk in TBST buffer (Tris-buffered saline with Tween 20, (50 mM Tris, 1.5 M NaCl, 0.1% Tween 20 (pH 7.4)) for 30 min at RT. Anti-KDEL (Enzo, ADI-SPA-827-F), anti-PDIA4 (ProteinTech, 14712−1-AP), anti-ASNS (ProteinTech, 14681-1-AP), and anti-GAPDH (GeneTex, GTX627408) antibodies were used to probe Western blots at a 1:1000 dilution in TBST blocking buffer (0.1% Tween, 5% BSA, 0.1% Sodium Azide).

### General Digestion Protocol (Manual and Automated)

#### Manual

20 µg of protein from each sample was aliquoted manually for label free DIA analysis. Proteins were precipitated using MS grade methanol (MeOH), chloroform, and water (H_2_O) in a 3:1:3 ratio and washed three times with 500 µL of MeOH. Each MeOH wash was followed by a 2 min centrifugation (21.1 kg, RT). Protein pellets were then air dried for 45 to 60 min and resuspended in 5 μL of 1% (w/v) Rapigest SF (Waters) in H_2_O. Resuspended proteins were subsequently diluted with 32.5 μL of water and 10 μL of 0.5 M HEPES (pH 8.0) and then reduced with 0.5 μL of freshly 0.5 M tris(2-carboxyethyl)phosphine (TCEP; Sigma) for 30 min at room temperature. Samples were then alkylated with 1 μL of fresh 0.5 M iodoacetamide (IAA; Sigma) for 30 min at room temperature in the dark and digested with 0.5 μg of Trypsin/Lys-C (Thermo Fisher; 1:40 protease to protein ratio) for 10 h.

### Automated

Protein cleanup and digestion was carried out on the Biomek i5 (Beckman Coulter). 20 µg of protein from each sample was aliquoted for label free DIA analysis or for TMT-DDA experiments using the automated sample hander. For both LC-MS/MS analysis workflows, proteins were digested on magnetic single pot solid phase enhanced sample preparation beads (SP3; Cytiva; 1:1 hydrophilic to hydrophobic beads) and cleaned-up as previously described^46^. Prior to digestion, proteins were reduced with fresh dithiothreitol (DTT; 5 mM, Sigma) for 30 min at 37°C and alkylated with fresh IAA (20 mM, Sigma) for 15 min at RT (in darkness). To quench the alkylation, DDT was added a second time (5 mM) with incubation for 15 min. With our protocol, protein was bound to SP3 beads with 100% ethanol and washed three times with 80% ethanol for clean-up. Proteins were subsequently digested with Trypsin/Lys-C (Thermo Fisher; 1:40 protease to protein ratio) in 50 µL ammonium bicarbonate (ABC; 100 mM; pH 8) or HEPES (0.5 M; pH 8) for 14 h (shaking at 700 rpm) for DIA and DDA-TMT analysis, respectively.

### Sample Preparation for LC-MS/MS after digestion

#### Manual Label Free DIA Sample Preparation

After digestion, formic acid (FA; ThermoFisher Scientific) was added to each sample to a final concentration of 4% (v/v, pH ∼2) with incubation at 37°C for 30 min to cleave Rapigest SF. Samples were subsequently spun down at 21.1 kg for 15 min and the supernatant was transferred to a fresh low-bind tube (ThermoFisher Scientific). Samples were subsequently dried *in vacuo* using a SpeedVac and stored dried at -80°C until use. Peptides were resuspended in 99 µL buffer A (4.9% ACN, 95% H_2_0, 0.1% FA (v/v/v). An additional 1 µL of FA was added prior to LC-MS/MS analysis for a final concentration of 200 ng of peptides per µL.

#### Biomek i5 Label Free DIA Sample Preparation

Digested peptides were removed from SP3 beads after digestion on the Biomek i5 and dispensed manually into low-bind tubes. FA was added to each sample to a final concentration of 2% (v/v) and samples were subsequently dried *in vacuo* using a SpeedVac (ThermoFisher Scientific, SPD111V). Samples were stored dried at -80°C until use. Peptides were resuspended in 99 µL buffer A and an additional 1 µL of FA. Samples were then spun down at 21.1k g for 15 min and then transferred to a fresh low bind tube prior to LC-MS/MS analysis.

#### Biomek i5 TMT Sample Preparation

Samples were labeled using 18-plex TMTpro (Thermo Scientific) reagents on the Biomek. To begin, the volume of each sample was raised to 60 µL with H_2_O. Peptides from each sample were then reacted with their respective tandem mass tag (resuspended in ACN) at 40% v/v for 1 h at RT. Reactions were then quenched by the addition of ABC (0.4% wt/v final concentration) and incubated at RT for 1 h. Lastly, samples were pooled and acidified with FA (added to reach pH 2). The sample was then concentrated on the SpeedVac to 1/3 of the pooled volume to remove ACN. The volume of the sample was then adjusted back to the original pooled volume with buffer A, centrifuged for 30 min at 21.1 kg to remove residual beads, and the supernatant was transferred to a fresh low bind tube prior to LC-MS/MS analysis.

#### LC-MS/MS Analysis

LC-MS/MS analysis was performed using an Exploris480 mass spectrometer (Thermo Fisher) equipped with a Dionex Ultimate 3000 RSLCnano system (Thermo Fisher). For both data acquisition methods, peptides were separated using a 21.5 cm fused silica microcapillary column (ID 100 μm) ending with a laser-pulled tip filled with Aqua C18, 3 μm, 100 Å resin (Phenomenex # 04A-4311). Electrospray ionization was performed directly from the analytical column by applying a voltage of 2.2 kV (positive ionization mode) with an MS inlet capillary temperature of 275 °C and a RF Lens of 40%.

For DIA analysis, 600 ng of peptides per sample were loaded onto a commercial trap column (C18, 5 μm, 0.3x5mm; ThermoFisher Scientific, 160454) using an autosampler. Peptides were then eluted and separated on a 2 h gradient with a constant flow rate of 500 nL/min: 2% B (5 min hold) was ramped to 35% B over 90 min and stepped to 80% in 5 min and held at 80% B for 5 min, followed by a 13 min hold at 4% B to re-equilibrate the column. Between injections, a 45 min column-wash was performed using the following gradient: 2% B (6 min hold) stepped to 5% over 2 min and subsequently ramped to a mobile phase concentration of 35% B over 7 min, ramped to 65% B over 5 min, held at 85% B for 8 min, then returned to 3% B for the remainder of the analysis.

The initial DIA method utilized in this study acquired MS/MS scans in a staggered window pattern with placements optimized by Thermo Fisher XCalibur instrument controls. Briefly, 61 x 20 *m/z* (390-1010 *m/z*) precursor isolation window MS^2^ DIA spectra (30,000 resolution, 1e6 automatic gain control (AGC) target, 55 ms maximum ion injection time (maxIT), 27% normalized collision energy (NCE)) were acquired within a single duty cycle to achieve an effective isolation window size of 10 *m/z*. A precursor spectrum (388-1012 m/z, 60,000 resolution, AGC target 3e6, 100 ms maxIT) was collected prior to the acquisition of both the 30 and 31 MS/MS scans across the mass range. Each scan cycle of 63 spectra took approximately 4.4 s.

Alternatively, to our initial DIA method our semi-staggered method consisted of 62 MS/MS scan events with a 10 *m/z* precursor isolation window and 1 *m/z* overlap (30,000 resolution, 3e6 normalized ACG target, 55 ms maxIT, 27% NCE, 40% RF lens) taken across a 380-1000 *m/z* mass range. A precursor spectrum (380-1000 *m/z*, 3e6 normalized ACG target, 40% RF lens) was taken at either 60,000 or 120,000 resolution with a maxIT of 110 or 240 ms, respectively. In both cases, loop control was utilized with n= 20 to intersperse MS^1^ scans between MS/MS acquisitions. The 60k and 120k resolution semi-staggered DIA methods comprised of 66 spectra were determined to have a duty cycle time of approximately 3.9 and 4.4 sec, respectively. All DIA data was collected in profile mode.

To perform TMT-DDA analysis, 10 µg of 18-plex TMT labeled peptides were loaded using a high-pressure chamber onto a triphasic MudPIT column (prepared as described previously).^22^ Briefly, the MudPIT column was packed with 1.5 cm of Aqua 5 µm C18 resin (Phenomenex # 04A-4299), 1.5 cm Luna 5 µm SCX resin (Phenomenex # 04A-4398), and 1.5 cm 5 µm C18 resin. Prior to analysis, the peptide loaded MudPIT column was desalted with buffer A for 30 min. Peptides were eluted from the first C18 phase onto the SCX resin using a 0.5 µL injection of buffer A with the following 90 min gradient: 4% B (5 min hold) ramped to a mobile phase concentration of 40% B over 35 min, ramped to 80% B over 15 min, held at 80% B for 5 min, then returned to 2% B in 5 min and then held at 4% B for the remainder of the analysis at a constant flow rate 500 nL/min. Peptides were then sequentially eluted from the SCX stationary phase using 10 µL injections of 10, 20, 30, 40, 50, 60, 70, 80, 90, and 100% buffer C (500 mM ammonium acetate in buffer A) in buffer A. The final fraction was eluted with 10% buffer B (99.9% acetonitrile, 0.1% formic acid v/v) and 90% buffer C. Fractions were collected using the following 140 min gradient with a constant flow rate of 500 nL/min: 4% B (10 min hold), ramped to 40% B over 90 min and stepped to 80% in 5 min and held 80% B for 5 min, returned to 4% B in 5 min and held for 25 min. To conclude the analysis and ensure all peptides are eluted from the last C18 section of the MudPIT column, 0.5 ul of buffer A was injected and eluted using the 140 min method described above. For both gradients, the loading pump was held at 5% B (5 µL/min) during the duration of the analysis. During buffer pickup and loading for each method, the loading pump was held at a flow rate of 10 µL/min for 10 min prior to the valve switch. 5 min prior to the end of the analytical gradient the valve was switched to the load position to prepare for sample loading and the flow rate was adjusted from 5 µL/min to 10 µL/min for the remainder of the analysis.

For TMT-DDA MS acquisition, a 3 s duty cycle was utilized consisting of a full scan (400-1600 *m/z*, 120,000 resolution) and subsequent MS/MS spectra collected in TopSpeed acquisition mode. For MS^1^ scans, the maximum injection time was set to 50 ms with a normalized AGC target of 100% and data was collected in profile mode. Only ions with an MS^1^ intensity above 1e4 with a charge state between 2-6 were selected for MS/MS fragmentation. The monoisotopic peak determination (MIPS) was set to peptide to ensure selection of peptides for fragmentation (relax restrictions when too few precursors are found was selected). Additionally, a dynamic exclusion time of 45 s was utilized (determined from peptide elution profiles) with a mass tolerance of +/- 10 ppm was used to maximize peptide identifications. MS/MS spectra were collected in centroid mode with a normalized HCD collision energy of 36, 0.4 m/z isolation window, automatic maximum injection time, a normalized AGC target of 100%, at a MS^2^ orbitrap resolution of 45,000 with a defined first mass of 110 *m/z* to ensure measurement of TMTpro reporter ions.

### Peptide Identification and Quantification

#### DIA

All DIA spectra were analyzed in DIA-NN (1.8.1) using a library-free search. A spectral library was generated using a Uniprot SwissProt canonical human FASTA database (downloaded 01May2024; containing 20,361 entries) and contaminant FASTA^47^ (containing 379 entries) included peptides 7 to 30 residues in length with one missed trypsin cleavage and a maximum of one variable modification (Met oxidation or N-terminal acetylation). N-terminal M excision was enabled and cysteine carbamidomethylation was included as a fixed modification. Precursors between *m/z* 380-1000 with charge states of 1-4 were considered. Additionally, fragment ions between *m/z* 200 and 1800 were included. Deep learning (using a single-pass mode neural network classifier) was then utilized to generate a new *in-silico* spectral library from the DIA data provided to then research the raw files against. Smart profiling was used for library generation. Unrelated runs (mass accuracy and scan windows determined separately for different runs), match between runs (MBR), heuristic protein inference, and no shared spectra were selected in the algorithm section of the DIA-NN GUI. Protein inference was made on genes and robust LC (high precision) was used as the quantification strategy. Protein assembly was performed in DIA-NN based on the parsimony principle. Lastly, retention time dependent cross-run normalization was performed within DIA-NN.

#### TMT-DDA

Peptide identifications and TMT-based protein quantification was performed using the Proteome Discoverer (PD) 2.4 (Thermo Fisher Scientific) platform. MS/MS spectra were searched using SEQUEST-HT against a Uniprot SwissProt canonical human FASTA database (downloaded 01May2024; containing 20,361 entries), contaminant FASTA^47^ (containing 379 entries), and a decoy database of reversed peptide sequences. Searches were carried out according to the following parameters: 20 ppm peptide precursor tolerance, 0.02 Da fragment mass tolerance, minimum peptide length of 6 AAs, trypsin cleavage with a maximum of two missed cleavages, dynamic modifications of Met oxidation (+ 15.995 Da), protein N-terminal Met loss (-131.040 Da), protein N-terminal acetylation (+ 42.011 Da), protein N-terminal Met loss and acetylation (+89.030), and static modifications of Cys carbamidomethylation (+57.021 Da) and TMTpro (N-terminus/ Lys, +304.207 Da). IDs were then filtered using Percolator to adjust the false discovery rate (FDR) at the peptide level to 1%. The Reporter Ion Quantification node in PD was used to quantify TMT reporter ion intensities. Only reporter ions with an average signal-to-noise ratio (S/N) greater than 10 to 1 (10:1) and a percent co-isolation less than 25 were utilized for quantification^48,49^. TMT reporter ion intensities were summed for peptides belonging to the same protein, including razor peptides. Proteins were filtered at 1% FDR and protein grouping was performed according to the parsimony principle.

#### Data Analysis (Proteomics)

Two custom R scripts, for both data acquisition types, were developed for data filtering, normalization, visualization, and UPR target filtering. To increase confidence in protein identifications, only proteins with two peptides per protein were retained for further analysis in data generated by both acquisition types. Additionally, contaminant proteins^47^ were filtered prior to subsequent analysis. No imputation was applied using either acquisition technique. Post generation of the normalized and log_2_ transformed protein matrices, individual fold changes were calculated per TMT channel (TMT-DIA) or injection (DIA) within R. The average fold change per treatment was subsequently calculated using our average.FC() function. The identified proteins in each dataset were then filtered against our branch-specific UPR target list to evaluate the state of the response. **DIA:** FDR (1%) was controlled by manually filtering precursor and protein q values present in the report file using our diann.dia.qc() function. Protein quantifications were calculated from precursor abundances normalized by retention time normalization in DIA-NN using the DIA-NN R package max_LFQ() function ^50^. Prior to fold change calculations (log_2_(Treatment/Control) and statistical calculations, DIA data was filtered at the protein level for a minimum number of observations using our filter.NA() function to ensure robustness of protein identifications. **TMT-DDA:** Global median normalization was performed using the medNorm() function with our R code. Using our function, correction factors per TMT channel were applied to the raw abundances of each observation in the corresponding channel. Statistical analyses were performed using GraphPad Prism software (version 10.0.2) or R (version 4.3.1). To determine which proteins were significantly differentially expressed, an unpaired two sample t-test with multiple testing corrections was utilized. Adjusted p values were calculated with the two-stage step-up method of Benjamini, Krieger, and Yekutieli. An adjusted *p* value less than 0.01 was considered statistically significant for all volcano plot visualizations. PCA analysis was computed on log_2_ transformed median normalized abundances, in which all NAs were imputed to one, using the prcomp package in R. For all analyses with three or more groups, a one-way ANOVA test with Tukey’s post hoc test was computed. A *p* value < 0.05 was considered statistically significant.

## Results

### Automated sample preparation and optimized DIA-MS method improve protein identification and reproducibility

To develop an optimized platform for UPR target protein detection, we first streamlined sample preparation and data-independent acquisition (DIA) methods. We took advantage of engineered stable cell lines (HEK293^DAX^ and Fv2e-PERK)^10,51^ that are inducible by small molecules to selectively activate the ATF6, IRE1/XBP1s or PERK branch of the UPR independently, allowing us to identify candidate branch-specific UPR targets (**Fig. 2A**). Briefly in the HEK293^DAX^ cell line, orthogonal small molecules trimethoprim (TMP) and doxycycline (DOX) selectively upregulate the abundance of either ATF6 and/ or XBP1s transcription factors utilizing destabilized domain (DD) and tetracycline (tet)-repressor technology, respectively^10^. Alternatively, the Fv2e-PERK HEK293 stable cell line expresses a chimeric protein consisting of the PERK cytosolic kinase domain fused to a dimerization domain that elicits a dose dependent response in the presence of a small molecule dimerizer (AP20187), resulting in Fv2e-PERK homodimerization.^52^ Proteins isolated from the activated stable cell lines were cleaned-up and digested using SP3 beads^46^ on the Biomek i5 automated sample handler (**Fig 2A**). To evaluate the peptide and protein identification performance of the automated sample preparation, we first compared manual and automated label free sample preparations. The automated sample preparations leveraging SP3 bead cleanup showed increases in protein identifications by 13-20% (4 sample preparation batches each with n=6, DMSO / TMP treated HEK293^DAX^ cells activating the ATF6 branch) compared to manual preparations using methanol/chloroform precipitation (**Fig. 2B**). A single preparation (n=6) was performed using the Biomek i5 with 10 µg of protein input and half the amount of SP3 beads as was used for 20 µg protein inputs. The reduction of protein input and beads did not result in a loss of protein identifications, indicating the method’s ability to accommodate samples with reduced input (**Fig. 2B**). Additionally, Biomek sample preparations showed low variability in protein identifications within sample batches, indicating the technique provides high reproducibility (**Fig. 2B**).

**Figure 2.**
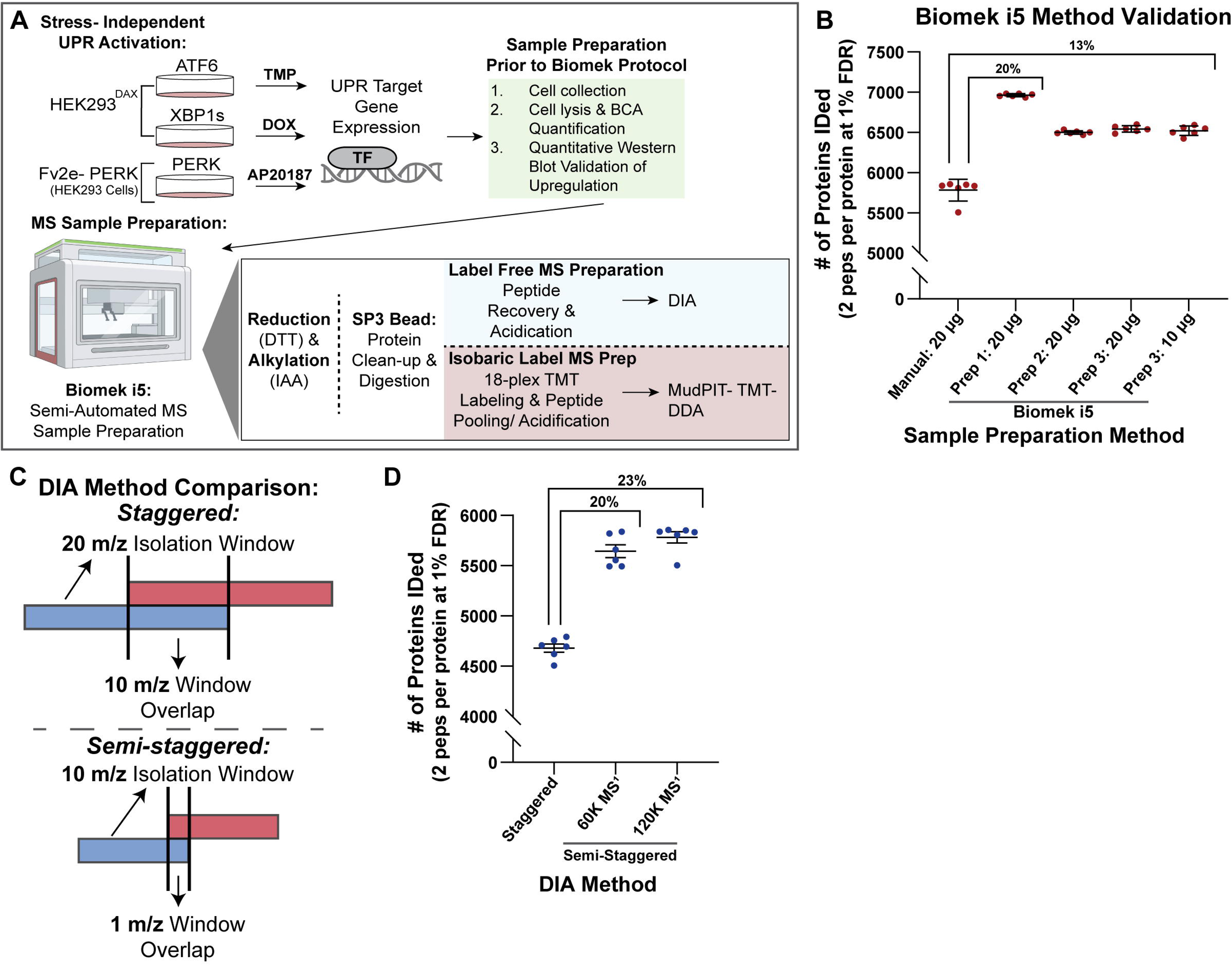
Validation of Biomek i5 label free sample preparation and DIA-MS methodologies. **(A)** Experimental schematic detailing the stable cell lines utilized for branch-specific UPR activation (HEK293^DAX^/ Fv2e-PERK), sample preparation on the Biomek i5 and the MS methods utilized in this study. Briefly, endogenous activation of ATF6 occurs when p90ATF6 translocates from the endoplasmic reticulum (ER) to the Golgi apparatus, where it is cleaved by S1P and S2P proteases, releasing the transcription factor p50ATF6. In the HEK293DAX engineered stable cell line, p50ATF6 is tagged with a destabilized DHFR (dDHFR) moiety, targeting it for degradation. However, treatment with TMP stabilizes the dDHFR, allowing p50ATF6 to function as a transcription factor. Alternatively, endogenous IRE1/XBP1s is activated when IRE1 autophosphorylates and non-canonically splices XBP1 mRNA into the active form, XBP1s, which then acts as a transcription factor. In HEK293^DAX^ cells, XBP1s expression is controlled by a doxycycline (dox)-inducible Tet-On system, allowing for direct transcription of XBP1s without the need for splicing. Lastly, endogenous PERK becomes active upon dimerization and autophosphorylation, which leads to phosphorylation of eIF2α. In the Fv2e-PERK cell line, activation is induced by a fusion protein containing the PERK kinase domain and two FKBP domains. Treatment with AP20187 promotes dimerization of the fusion protein, thereby activating PERK. Pharmacological activation of ATF6, XBP1s, and PERK was achieved with TMP (10 µM), doxycycline (1 µg/mL), and AP20187 (5 nM) for 16-18 hours, respectively. (**B**) The number of proteins identified (filtered at two peptides per protein at 1% FDR) between a manual (Methanol/ Chloroform precipitation) and three different automated label free sample preparations (Biomek i5, SP3 clean-up/ on-bead digestion) (n=6) analyzed by DIA-MS (120K MS^1^ revised method). (**C**) Schematic showing the DIA methods compared in this study, staggered versus semi-staggered (MS^1^ resolution of 60K or 120K). Methods differ in their isolation window size and degree of window overlap. The semi-staggered DIA method (120K MS^1^ resolution) was utilized for comparisons to TMT-DDA and to determine new branch specific UPR targets in subsequent figures. (**D**) Scatter dot plot depicting the number of identified proteins (filtered at two peptides per protein at 1% FDR) in the manually prepared samples (n=6). Both semi-staggered DIA methods (MS^1^ scans taken at a resolution of either 60 k or 120 k) showed a 20-23% increase in proteins identified compared to the initial DIA method.

Additionally, to increase the number of protein identifications attainable using DIA on our Exploris 480 mass spectrometer, we evaluated two different acquisition approaches. The initial method followed a conventional wide isolation window of 20 *m/z* with 10 *m/z* window overlap (*staggered acquisition*; **Fig. 2C**). In the enhanced method, the size of the MS^2^ isolation window and the MS^2^ window overlap were reduced to 10 *m/z* and 1 *m/z (semi-staggered acquisition),* respectively (**Fig. 2C**). Loop control was added in the new method to intersperse a MS^1^ scan after every twenty MS^2^ scans. We also compared a MS^1^ resolution of 60k and 120k. To divulge the utility of the altered DIA methods, we analyzed peptides prepared manually from HEK293^DAX^ cells (DMSO and TMP treated) and compared overall protein identifications and reproducibility between methods. The updated DIA methods at either MS^1^ resolution showed increases in identified precursors (38-40% at 1% FDR) and identified proteins (20-23%, 2 peptides per protein at 1% FDR) (**Fig. 2D**). These methods were determined to be reproducible without reductions in quantification accuracy (**Supplemental Fig. S1**). Therefore we utilized the DIA method with a MS^1^ resolution of 120k (semi-staggered window) throughout this study with spectral interpretation and quantification using DIA-NN (version 1.8.1)^53^.

### DIA-MS outperforms TMT-DDA-MS in the detection of UPR target proteins

We next sought to compare UPR target protein detection capabilities of two MS acquisition techniques, DIA-MS and TMT-DDA-MS. Many proteostasis related proteins regulated by the UPR are low abundant, making them challenging to quantify due to lack of detection of their associated peptides. To determine which method most comprehensively identifies and quantifies UPR targets, the HEK293^DAX^ and Fv2e-PERK cell lines were again utilized. Activation of the ATF6, XBP1s, PERK branches of the UPR were validated in HEK293^DAX^ and Fv2e-PERK samples via quantitative western blot (**Supplemental Fig 2 and 3**).

**Figure 3.**
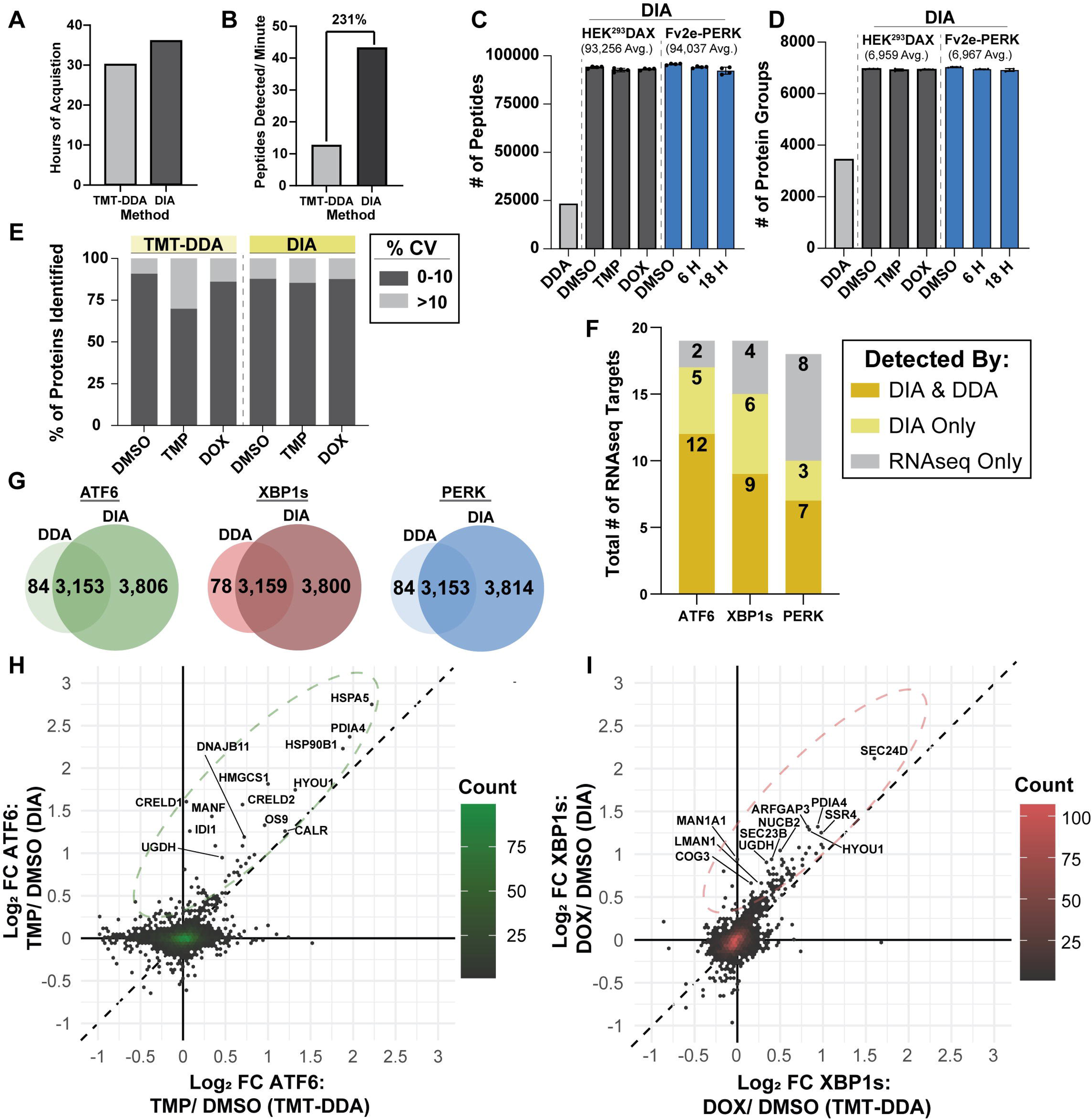
Comparing MS acquisition methods (DIA vs. TMT-DDA) to measure UPR target proteins. **(A)** Time required to measure 18 samples by either TMT-DDA (18-plex TMTPro with 12 fractions) or DIA (unfractionated). **(B)** Bar graph comparing the number of peptides detected per minute in the two acquisition methods. Bar graphs comparing the number of peptides **(C)** and protein groups **(D)** detected in the two stable cell lines when treated with their respective activators. **(E)** Percentage of proteins identified with a coefficient of variation (% CV) of between zero and 10 percent and greater than 10 percent in HEK^293^DAX cells treated with DMSO (0.1% v/v, TMP (10 µM), and DOX (1 µg/ mL) and measured with either TMT-DDA or DIA. **(F)** Stacked bar graph indicating the number of branch-specific UPR target genes previously described for RNAseq analysis^6^ that were measured by MS (TMT-DDA and DIA). **(G)** Venn diagrams indicating the degree of overlap in protein identifications between DDA and DIA in ATF6, XBP1s, and PERK activated cells. DIA was able to identify approximately 97% of the proteins identified via DDA. **(H-I)** Correlation plots depicting the log_2_ fold changes (FC) of proteins measured via TMT-DDA and DIA in the HEK293^DAX^ engineered stable cell line with either **(H)** ATF6 or **(I)** XBP1s activated. Colored dashed lines ovals indicate proteins that have a higher fold change in DIA compared to TMT-DDA. The black dashed line indicates y=x.

For the preparation of TMT labeled peptides, we used a previously developed and validated Biomek i5 TMT labeling protocol^54^. Upon comparison with our manual TMT preparations, the automated method resulted in a similar number of peptide and protein identifications. Owing to the improved identifications observed from our Biomek method, all samples in this study were prepared using the automated sample handler. The time required to acquire 18 samples on the DIA method utilized throughout this study was slightly longer than the TMT-DDA method, at 36 and 30 hours respectively (**Fig. 3A**). However, DIA analysis significantly outperformed DDA, detecting approximately 231% more peptides per minute (**Fig. 3B**). Similarly, DIA analysis showed a 304% average increase in peptide group detection, identifying 93,521 peptide groups compared to 23,154 detected by TMT-DDA (**Fig. 3C**). This increase in detected peptides was also reflected in the number of protein groups identified, with DIA detecting 6,953 protein groups versus 3,443 detected by TMT-DDA (**Fig. 3D; Supplemental Table S2, S3, S4**).

To compare the precision of DIA and TMT-DDA based quantification, the percent coefficient of variation (% CV) of proteins identified in ATF6 and XBP1s activated HEK293^DAX^ cells was determined (**Fig. 3E)**. Both methods showed a similar percentage of proteins with a % CV greater than 10% across treatments. Overall, these findings indicate that both DIA and TMT-DDA methods offer comparable quantification precision at the protein level, regardless of sample multiplexing when using TMT. Additionally, 3,237 of the 3,443 proteins identified across TMT-DDA were quantifiable. This means that approximately 6% (206 proteins) were not quantifiable due to a signal-to-noise ratio of less than 10:1 and/or co-isolation interference greater than 25%. As expected, DIA resulted in an incomplete protein matrix, likely due to non-stochastic missing values within the matrix of 6,953 protein groups^55^. However, per injection in the DIA protein matrix only approximately 3% of proteins were not detected and/ or not quantifiable resulting in an “NA” designation, which was two-fold less than TMT-DDA. To provide higher confidence in protein presence in DIA, the protein matrix was filtered for a minimum of three of four observations per treatment per protein prior to statistical analysis.

Importantly, we identified a greater number UPR-branch specific genes targets (as initially defined from RNAseq – **Fig. 1B**) in our DIA dataset compared to TMT-DDA^6^ (**Fig. 3F**), confirming the improved sensitivity of the DIA-MS workflow. Regardless of the treatment applied to induce branch-specific UPR activation, DIA was able to detect approximately 97.5% of the proteins identified in the TMT-DDA dataset (**Fig 3G**), revealing a high degree of overlap in the proteins identified by both techniques. Despite the increase in sensitivity afforded by DIA acquisition, we were unable to detect the master regulators (e.g. XBP1s, PERK, CHOP, GADD34, etc) of the UPR, except for ATF6. However, these proteins are very low in abundance and have previously required an optimized parallel reaction monitoring (PRM) method on an orbitrap mass spectrometer for detection^56^.

The associated log_2_ fold changes of proteins detected by DIA were generally greater than those observed via TMT-DDA analysis. In ATF6 activated stable cell lines, fold changes were approximately 1.3 times greater in DIA measurements compared to DDA, indicated by the green dashed oval on the correlation plot (**Fig. 3H**). Similarly, XBP1s activated cell line DIA measurements were roughly 1.2 times greater than DDA (red dashed oval; **Fig. 3I**). The superior ability of DIA to identify a greater number of UPR regulated proteins prompted the use of this technique for the branch-specific classification of UPR regulated proteins and identification of new branch-specific UPR targets.

### Correlation analysis of ATF6 and XBP1s engineered stable cell line proteomes reveals new branch-specific UPR markers

To identify proteins previously undescribed as regulated by the UPR and subsequently classify them to a specific UPR branch, we performed an unbiased correlation analysis (**Fig. 4A**). We first compared ATF6 and XBP1s branches since the majority of genes activated by ATF6 are also activated by XBP1s at varying degrees^10^. Proteins with a log_2_ fold change greater than 0.5 ± 0.05 towards either the ATF6 or XBP1s branch were designated as branch-specific targets (**Fig 4A; Supplemental Table S5, S6, S7, S8**). The classified proteins were then further validated for regulation by the UPR by cross-referencing for significant upregulation (1% FDR) in multiple datasets. First, two DIA proteomics datasets with global UPR activation were generated by treating HEK293T cells with ER stressors thapsigargin (Tg; 500 nM; 16 h) and tunicamycin (Tm; 500 nM; 16 h) (**Supplemental Fig. S4; Supplemental Table S9**). Tg and Tm globally activate the UPR by causing accumulation of misfolded proteins in the ER lumen through distinct mechanisms. Tg depletes ER luminal calcium stores through inhibition of the sarcoplasmic/ endoplasmic reticulum Ca^+2^ ATPase (SERCA), impairing calcium-dependent chaperones^57^, while Tm inhibits N-linked glycosylation resulting in an accumulation of unfolded glycoproteins in the ER^58^. Activation of select UPR markers in Tg and Tm treated cells was first validated using quantitative western blot analysis (**Supplemental Fig. 5**). Second, we mined existing RNAseq dataset (Tg treated for 16 h)^59^. ATF6 candidate proteins were further referenced against an additional RNAseq dataset of HEK293^DAX^ cells treated with TMP (**Fig. 4A**)^59^.

**Figure 4.**
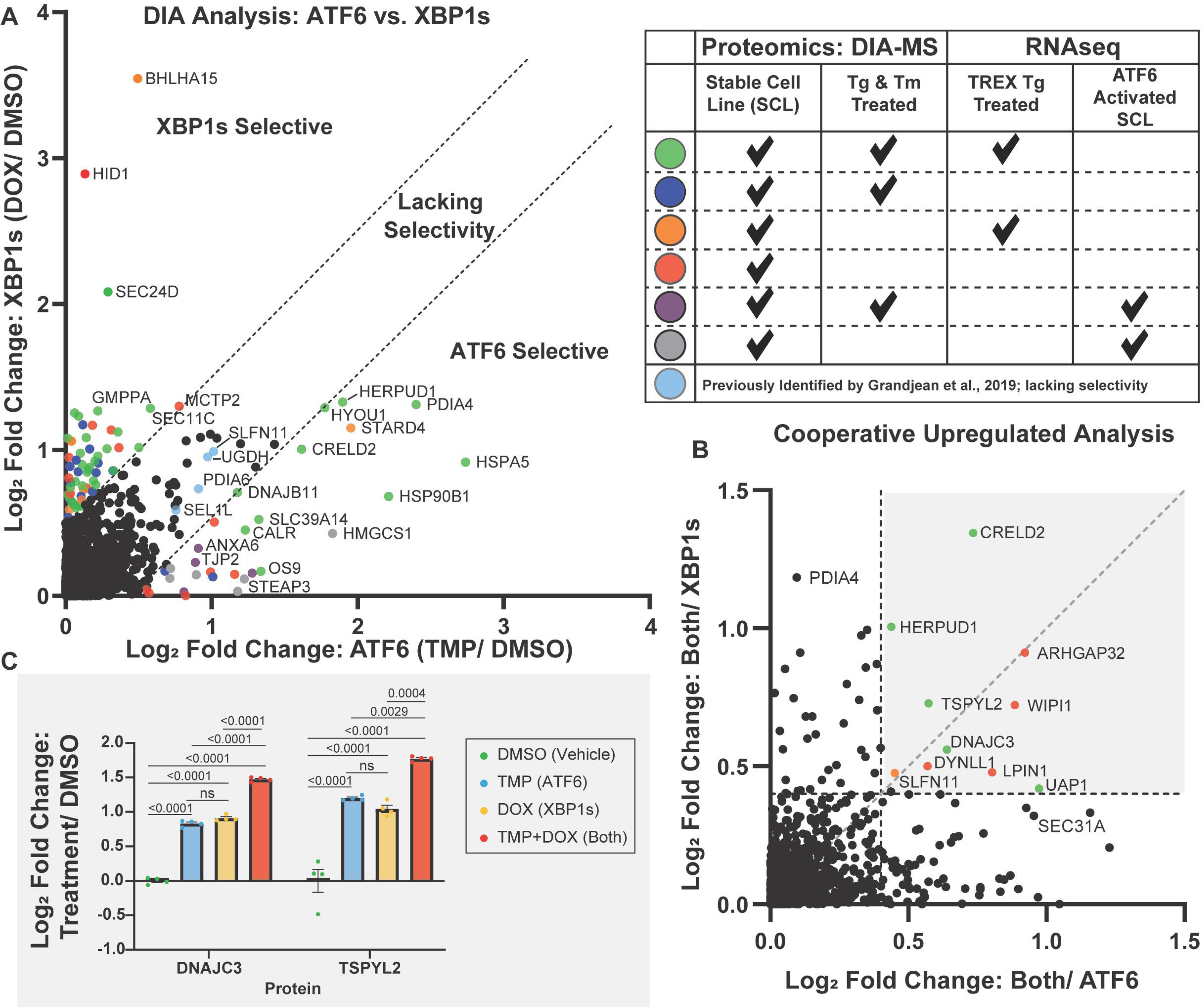
Determination of ATF6 and XBP1s branch-specific proteomics targets and cooperatively regulated proteins with HEK293^DAX^ cells. **(A)** Correlation plot of the log_2_ fold change of ATF6 (x-axis, TMP/DMSO) and XBP1s (y-axis, DOX/DMSO) in activated HEK293^DAX^ engineered stable cell lines measured via DIA-MS. Proteins were categorized into three categories: ATF6 selective, XBP1s selective, or lacking specificity based on the log_2_ fold change cutoff of 0.5 ± 0.05 as indicated by dotted lines. Each dot represents an individual protein with colors representative of whether or not the protein is present in global UPR activated proteomics DIA datasets (16 h treatment of HEK293 cells with Tg (500 nM) or Tm (500 nM)) and/ or RNAseq datasets. Only proteins with a green dot were selected to be branch-specific UPR protein targets (highest confidence). **(B)** Correlation analysis to identify cooperatively upregulated UPR proteins. The black diagonal dashed line indicates y=x. Vertical and horizontal lines indicate a log_2_ fold change greater than 0.4. **(C)** Bar plots of two newly identified cooperatively upregulated UPR target proteins, DNAJC3 and TSPYL2, quantified via DIA-MS. n=4, mean ± SEM. Statistical analysis was performed using one-way ANOVA per protein with Benjamini, Krieger, and Yekutieli multiple testing correction, p <0.05 considered statistically significant.

To curate a robust list of branch-specific UPR protein targets, emphasis was placed on genes which are significantly upregulated at both transcriptional and translational levels to provide greater confidence of UPR regulation. To this end, we removed proteins that were either absent or not significantly upregulated in either Tg or Tm-activated DIA-MS proteomics datasets and/ or the RNAseq reference dataset (TMEM50B, MIS12, SLFN11, STARD4, PLEKHA6, etc.). Lack of detection of proteins in the DIA-MS dataset may be due to peptide interference resulting in removal during data processing^53^, inefficient isolation during cell lysis, or absence in the chronic UPR translational response compared to the acute response. Several genes previously identified by Grandjean et al. as ATF6-specific were upregulated to a similar degree in the DAX cell lines when XBP1s was induced (e.g. MANF, PDIA6, SEL1L, and UGDH). Since these genes lack ATF6 or XBP1s selectivity, they were omitted from our branch-specific proteomics target list. However, since many of these genes were significantly upregulated in RNAseq and Tg/Tm datasets they were labeled as “UPR Regulated - Not Branch Specific” (**Supplemental Table S10**).

ATF6 and XBP1s transcription factors can heterodimerize, resulting in a unique transcriptional and translational profile^10,11^. Cooperative upregulation thus results from combined XBP1s and ATF6 activation as a result of either the binding of both XBP1s and ATF6 to promoter regions or the preferential binding of XBP1s/ATF6 heterodimers to select promoters. In this case, genes are upregulated to higher levels than when activating either XBP1s or ATF6 alone. To interrogate cooperativity in the HEK293^DAX^ stable cell line, we concurrently treated cells with TMP and DOX, to upregulate ATF6 and XBP1s transcription factors, respectively. To identify proteins whose expression is regulated cooperatively, we generated a correlation plot comparing concurrent treatment against selective activation of ATF6 or XBP1s (**Fig. 4B**). Proteins closest to the y=x line (gray) show the strongest cooperative upregulation whereas proteins close to the axes indicate preferential dependency on either transcription factor. As determined previously from Shoulders et al., CRELD2 is induced cooperatively^10^. Importantly, several other proteins were cooperatively regulated by XBP1s and ATF6 (CRELD2, DNAJC3, HERPUD1, TSPYL2, and UAP1), which were subsequently validated using our global UPR DIA datasets and RNAseq. Colors of each dot, representing a protein, indicate in which datasets (global UPR activated or RNAseq) each protein was also identified for validation. Five of the ten proteins identified as cooperatively regulated through the correlation analysis after validation with additional datasets (**Fig. 4C**). These cooperatively regulated proteins were removed from branch-specific target protein lists and labeled as “cooperatively regulated” (**Supplemental Table S10**).

### Defining ATF6 and XBP1s Branch-Specific Proteomics Targets

We compiled a final list of ATF6 and XBP1s branch-specific proteins identified in this study (**Fig. 5A**). In addition to observing eleven genes previously described as XBP1s branch-specific transcriptomics targets in our proteomics datasets, our approach identified an additional 23 branch-specific XBP1s proteomics targets. Of the genes previously described by Grandjean et al., 2019 as being regulated by XBP1s transcriptionally, four of these genes (i.e. EDEM2, PLPP5, FICD, and MBNL2) were not identified in proteomics datasets and/or were not identified or not significantly enriched (1% FDR) in the RNAseq dataset utilized for analysis. Additionally, numerous genes were removed from the original list as they were not detected or significantly upregulated in Tg and Tm proteomics datasets (i.e. DNAJB9, STT3A, SRP19), highlighting the discrepancies between branch-specific transcriptomics and proteomics targets for the UPR. The list of branch-specific ATF6 markers was reduced compared to transcriptomics, largely because ATF6/XBP1s-cooperatively regulated genes were removed. The log_2_ fold change of the branch-specific UPR proteins were calculated from Tm and Tg UPR activated datasets and were plotted as correlation plots for ATF6 (**Fig. 5B; Supplemental Fig. 6A/B)** and XBP1s (**Fig. 5C; Supplemental Fig. 6A/B).** Activation with Tg and Tm resulted in the differential upregulation of some XBP1s and ATF6 targets. For example, HYOU1, PDIA4, and SEC24D showed a significant difference in their degree of regulation when treated with Tm or Tg (**Fig. 6A/B**). Overall, these data indicate that the nature of the cause of UPR activation impacts the degree of activation of a particular UPR regulated protein^60^ (**Supplemental Fig. 4G/H**).

**Figure 5.**
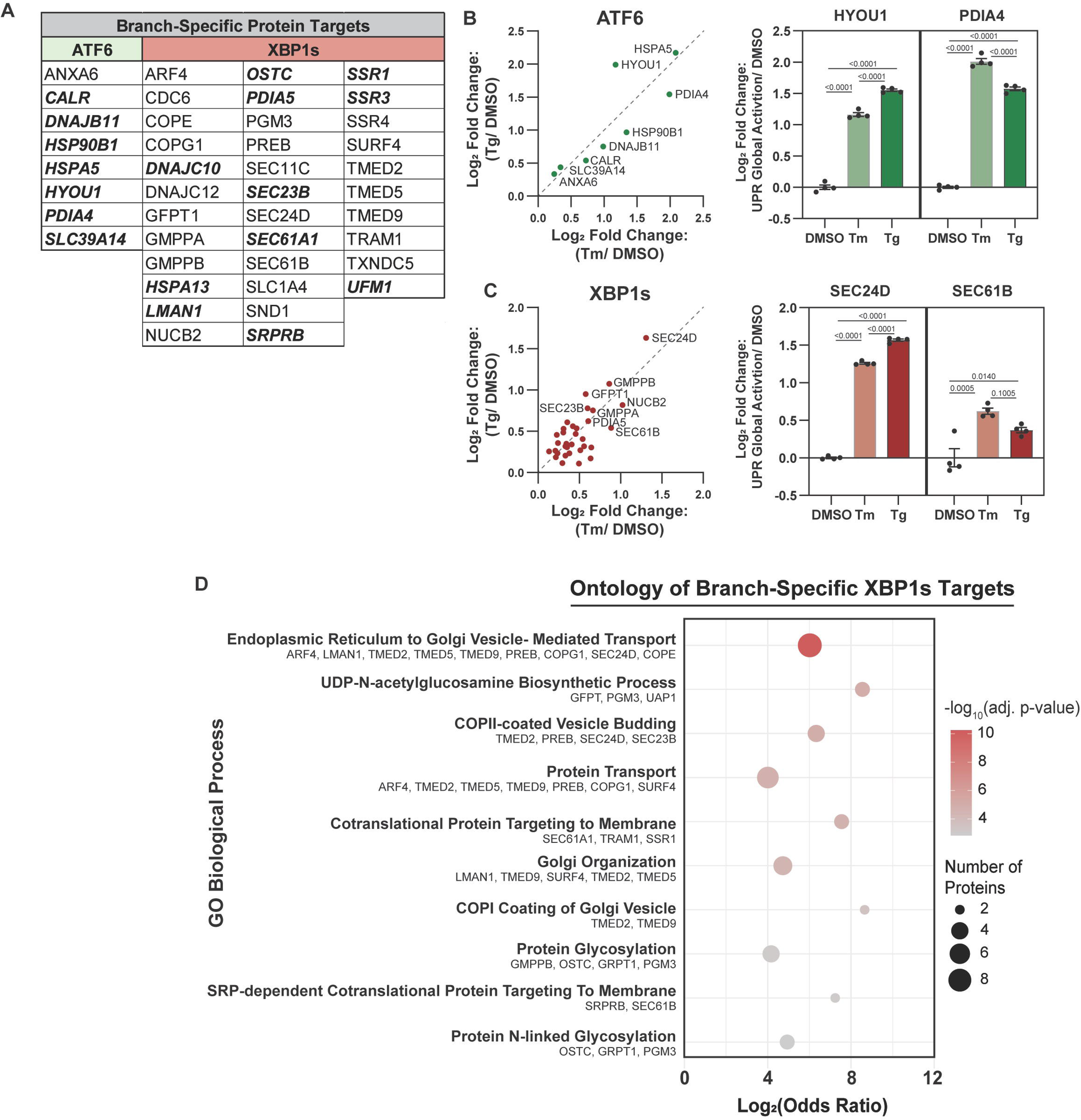
Validation of ATF6 and XBP1s branch-specific proteomics targets. **(A)** List of the ATF6 and XBP1s branch-specific UPR proteins determined in this study. ***Bold/ italized*** proteins were present in Grandjean et al., 2019 RNAseq analysis. Correlation plots of the log_2_ fold change of proteomics UPR targets shown in (A), for ATF6 (**B**) and XBP1s (**C**), measured in HEK293 cells activated with either Tm or Tg compared to vehicle (DMSO). Quantification of select targets plotted as a bar graph for each global UPR treatment. Error bars represent the mean ± SEM; (n =4). Statistical analysis was performed using one-way ANOVA with Benjamini, Krieger, and Yekutieli multiple testing correction, p <0.05 considered statistically significant. **(C)** Gene ontology of the XBP1s protein targets established in this study, shown in (A), using Enrichr^88^. Adjusted p-value was computed using the Benjamini-Hochberg method to correct multiple hypothesis testing.

**Figure 6.**
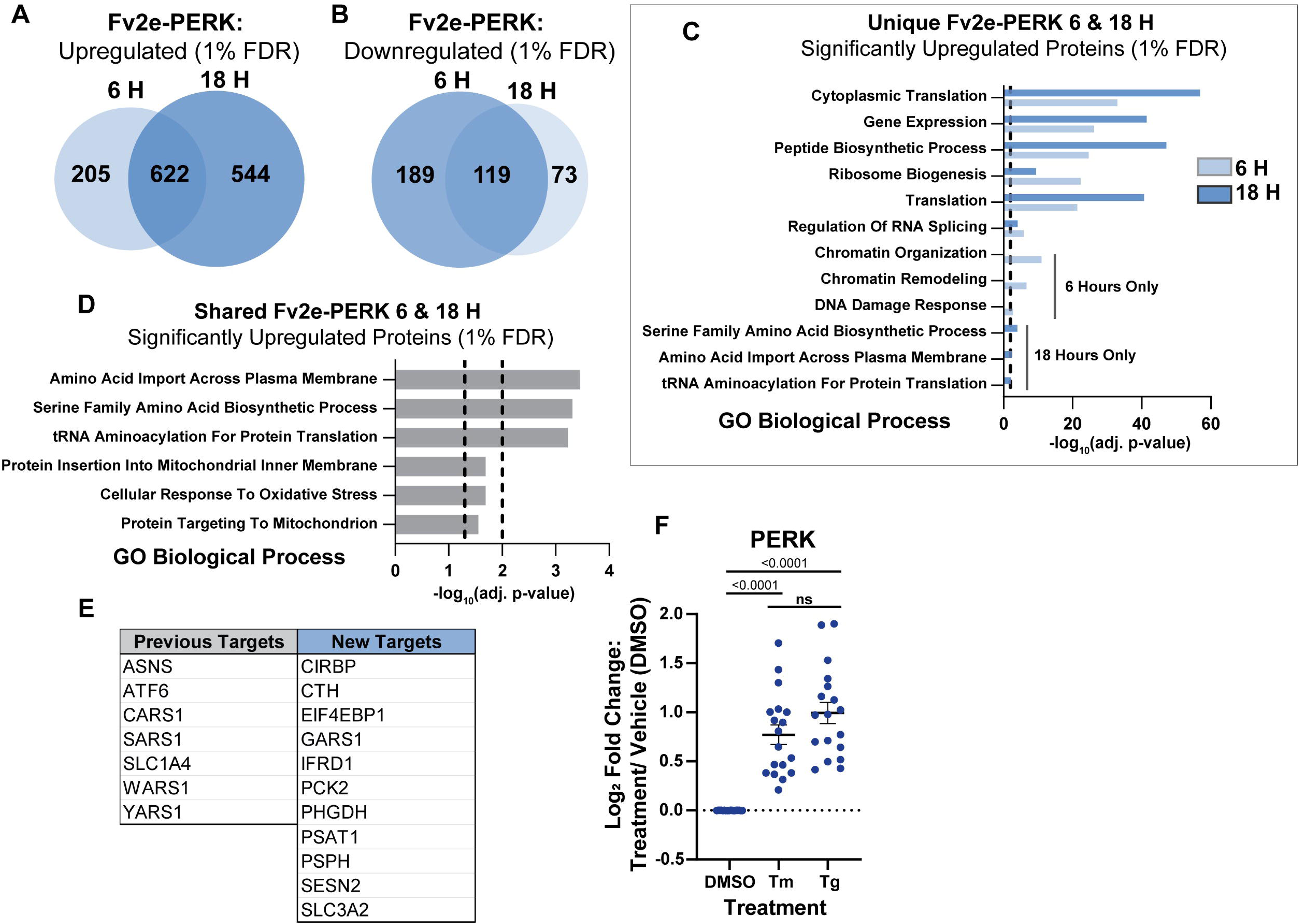
Determination of PERK branch-specific proteomics targets using Fv2e-PERK cells. (**A-B**) Venn diagrams to visualize the number of downregulated (**A**) and upregulated (**B**) proteins that are shared and unique in Fv2e-PERK stable cells 6 and 18 hours following PERK activation with AP20187 (5 nM). (**C**) Gene ontology of proteins uniquely upregulated at 6 and 18 hours (not including shared proteins) in Fv2e-PERK stable cells using Enricher. Adjusted p-value was computed using the Benjamini-Hochberg method for correction for multiple hypothesis testing. (**D**) Gene ontology (using Enricher) significantly enriched proteins (1% FDR) shared between Fv2e-PERK cells treated with AP20187 (5 nM) for either 6 or 18 hours. The black lines indicate a -log_10_(adjusted p-value) of 1.3 and 2. (**E**) List of the PERK branch-specific UPR proteins determined in this study after proteomics validation. Previous targets include those retained from Grandjean et al., 2019 (RNAseq analysis). (**F**) Proteomics UPR targets from table (E) for PERK activation detection measured in HEK293 cells activated with either Tm or Tg (500 nM; 16 h). Statistical analysis was performed using one-way ANOVA with Benjamini, Krieger, and Yekutieli multiple testing correction, p <0.05 considered statistically significant.

ATF6 targets are largely chaperones or co-chaperones, as previously described by Shoulders and colleagues^10^. Gene ontology analysis of proteins identified as XBP1s branch-specific revealed involvement in ER to Golgi vesicle-mediated transport (both COPI and COPII trafficking), protein glycosylation, UDP-N-acetylglucosamine (UDP-GlcNAc) biosynthesis, and ER associated degradation (ERAD) (**Fig. 5C**). The hexosamine biosynthetic core pathway (HBP) produces UDP-GlcNAc, a metabolite used for N- or O-linked glycosylation. Here, we see significant upregulation of the key enzymes: GFPT1, PGM3, and UAP1, with UAP1 being cooperatively regulated (**Supplemental Fig. 6B**). Our dataset specifically ascribes this regulation of the HBP to the IRE1/XBP1s branch. Additionally, proteins from Coat Protein I (COPI) and Coat Protein II (COPII) families were found to be induced by XBP1s. Upregulated COP1 machinery included subunits of the COPI complex such as COPG1 and COPE, whereas COPII associated proteins included SEC24D, SEC22B, and PREB. Secondly, TMED proteins (TMED2, TMED5, TMED9), known to be involved in anterograde transport, were also found to be significantly upregulated by XBP1s^61–63^.

### Time-dependent analysis of the Fv2e-PERK engineered stable cell line to determine branch-specific PERK target proteins

To identify branch-specific PERK proteomic targets, we compared the proteomes of Fv2e-PERK cells treated with AP20187 for either 6 or 18 hours against DMSO vehicle to identify significantly upregulated and downregulated proteins (1% FDR) (**Supplemental Fig. 7; Supplemental Table S11, S12, S13**). The PERK response, known for its dynamic nature, involves translational repression that triggers a positive feedback loop to restore translation, highlighting the importance of evaluating the proteome at multiple timepoints ^64^. PERK activation results in eIF2α phosphorylation which induces transient global translation attenuation to reduce the flux of proteins entering the ER for folding^4,12^. Western blotting confirmed that a greater proportion of eIF2α was phosphorylated at 6 hours compared to 18 hours, as expected (**Supplemental Fig. 3**). In the proteomics dataset, the number of significantly upregulated proteins was greater at 18 hours post PERK activation compared to 6 hours, although there was an extensive overlap between both timepoints (**Fig. 6A**). Interestingly, a greater number of proteins were downregulated at 18 hours as opposed to 6 hours (**Fig. 6B**). These data point to the temporal nature of the PERK response. To investigate the biological processes which differ between the two timepoints, gene ontology was carried out using Enrichr on significantly upregulated proteins unique to the 6- or 18-hour collection timepoints (**Fig. 6C**). At both timepoints, several biological processes, such as cytoplasmic translation and ribosome biogenesis, exhibited upregulation. However, distinct sets of proteins were involved in mediating these biological functions at the two timepoints studied. Proteins involved in chromatin remodeling and DNA damage response were particularly expressed at 6 hours. Specifically at 18 hours, there is an upregulation of amino acyl-transferases and amino acid transporters. These unique profiles at the two time points indicate the progression of the response can be monitored by proteomics. Additionally, our Fv2e-PERK proteomics dataset showed significant enrichment (1% FDR) of proteins shared between the two timepoints (6 and 18 hours) involved in response to oxidative stress, mitochondrial processes and amino acid synthesis/ transport (**Fig. 6D**).

To identify protein targets indicative of chronic PERK activation, we considered significantly upregulated (1% FDR) proteins observed at the 18-hour timepoint in Fv2e-PERK cells. We validated the identified proteins using our Tg and Tm (16 hour) treated global DIA proteomics datasets and a Tg treated (16 hour) RNAseq dataset. Proteins which were significantly upregulated (1% FDR) in the three confirmation datasets were deemed UPR regulated and were combined with validated PERK/ISR RNAseq targets previously identified by Grandjean et al. (**Fig. 6E**). Collectively, the target PERK proteins were significantly upregulated in cells treated with global UPR activators for extended time periods (16 hours) (**Fig. 6F**).

### BRAF-mutant Melanoma Cell Lines Show Differential Branch-Specific UPR Activation

Activation of the UPR in response to ER stress has been previously described in the survival of melanoma cells to accommodate increased demands for protein and lipid production in rapidly dividing cells^65–67^. Drug-tolerant idling BRAF-mutant melanoma cells possess frequent spontaneous [Ca^2+^]_cyt_ (cytosolic) signals in up to 60% of the population^34^. Further investigation revealed ER Ca^2+^ stores were reduced, potentially due to abundant [Ca^2+^]_cyt_ signals driving a high rate of Ca^2+^ efflux^34^. However, since Ca^2+^ is critical in the maintenance of ER homeostasis, Ca^2+^ depletion is a potent inducer of ER stress^68^ and may be activated here.

To determine if the UPR is activated in the drug-tolerant idling cells with aberrant calcium signaling, we evaluated the response using our branch-specific UPR protein markers after 3 and 8 days of treatment with PLX4720, an analog of the clinical BRAF inhibitor vemurafenib^43,44^. Four separate patient-derived cell lines harboring the activating BRAF-V600E mutation (A375, SKMEL5, SKMEL28, WM88)^33,69^ were investigated (**Fig. 7A**). Cells were collected and analyzed via DIA LC-MS/MS at 3 days and 8 days with treatment (**Supplemental Fig. 8A, 8B, Supplemental Table S13**). Two timepoints were compared since previous data revealed calcium spiking activity became more prevalent across the population with increasing BRAFi treatment time, indicating duration of exposure to PLX4720 can influence the non-genetic mechanisms at play in drug tolerance^34^.

**Figure 7.**
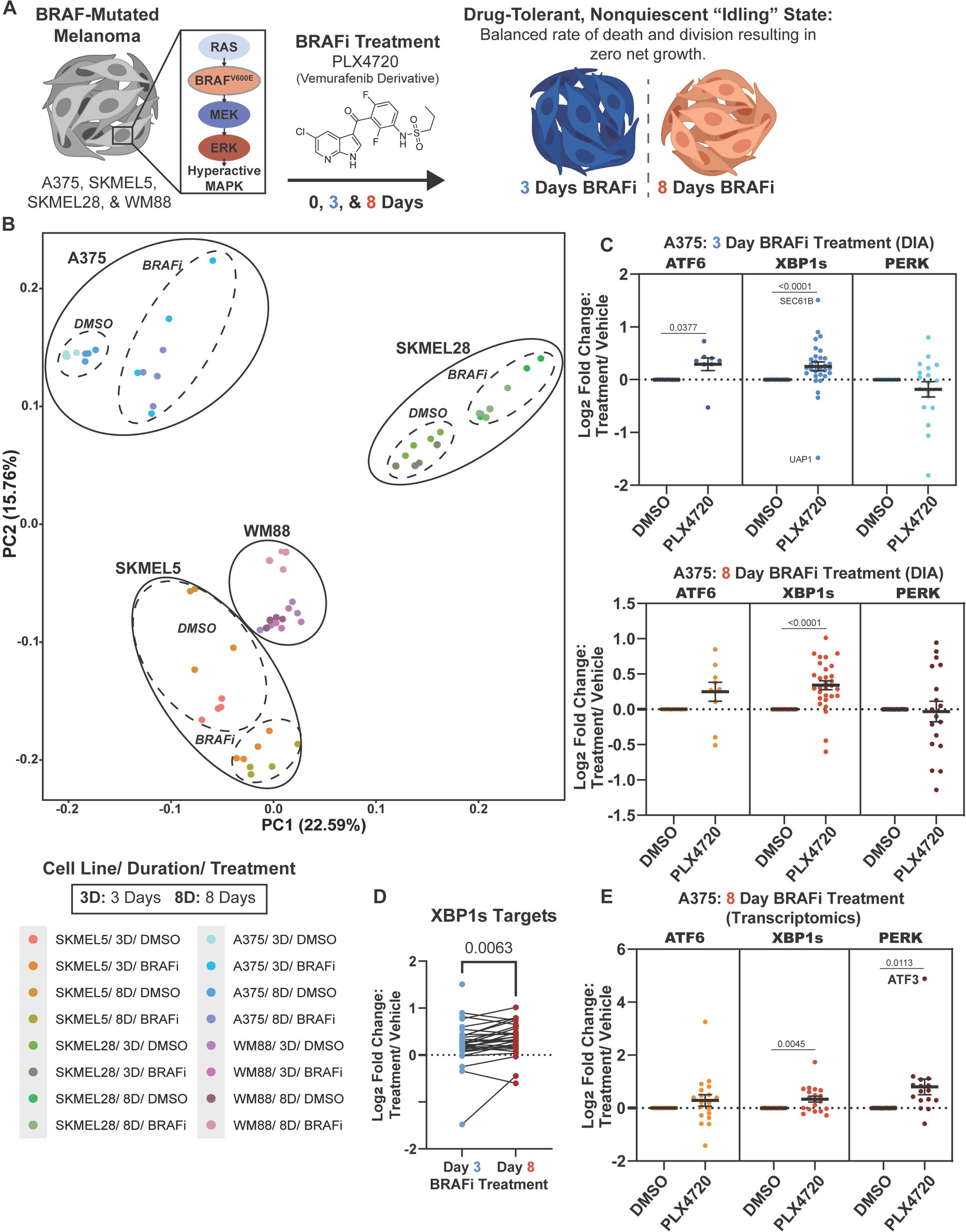
Evaluation of branch-specific UPR activation in A375 cells using proteomics. **(A)** Schematic overview of the BRAF-mutant melanoma cell lines and treatment (including durations) utilized in this study. BRAF is part of the MAP kinase pathways and cell lines contain the V600E mutation. Cells were treated with the BRAF inhibitor PLX4720, an analog of Vemurafenib. **(B)** Principal component analysis (PCA) of LC-MS/MS proteomics data collected on four BRAF-mutant melanoma cell lines (SKMEL5, SKMEL28, A375, and WM88) treated for 3 and 8 days with PLX4720 or vehicle (DMSO). **(C)** Measurement of the UPR protein targets for all three branches of the UPR, in A375 cells, after 3 and 8 days with PLX4720 treatment. **(D)** Comparison of the log_2_ fold change of XBP1s protein targets after 3 and 8 days of PLX4720 treatment. (**E**) Measurement of the UPR transcriptional targets for all three branches of the UPR after 8 days with BRAFi treatment in A375 cells from published RNA-seq data^34^. Statistical analysis was performed using an unpaired t-test, p <0.05 considered statistically significant.

PCA analysis of all samples revealed a clear separation between the cell lines, as well as between the PLX4720 treatment compared to vehicle (DMSO). In contrast, less separation of the clusters was observed in cells treated for 3 or 8 days (**Fig. 7B**). Proteomics analysis using DIA, combined with our branch-specific target protein sets, identified activation of ATF6 and XBP1s in the A375 cell line after 3 days and both 3 and 8 days of PLX4720 treatment, respectively (**Fig. 7C**). The abundance of the XBP1s target proteins increased significantly between days 3 and 8 (**Fig. 7D**). To further corroborate the XBP1s activation, a previously published bulk RNAseq data^34^ was analyzed using the branch-specific UPR transcriptomics targets described by Grandjean and colleagues. XBP1s activation was additionally observed in the transcriptomics analysis of the A375 cell line after 8 days of PLX4720 treatment. (**Fig. 7E**)^6^. Protein targets of the PERK branch were not found to be differentially expressed at either timepoint in the proteomics dataset (**Fig. 7C**). However, transcriptomics analysis indicated activation of the PERK branch (**Fig. 7D**). Alternatively, while the ATF6 branch of the UPR was not found to be significantly upregulated in aggregate, many of the protein markers used to assess this branch of the pathway were upregulated (**Fig 7C**). Similarly to the A375 cell line, the SKMEL28 cell line additionally showed significant XBP1s activation with 8 days of BRAFi treatment. Alternatively, XBP1s targets in aggregate appeared unchanged or downregulated in WM88 and SKMEL5 cells with 3 and 8 days of BRAFi treatment, respectively (**Supplemental Fig. 9**). Through the combination of our validated proteomics targets and DIA, we established a high-throughput approach to dissect branch-specific UPR activation in drug-tolerant BRAF-mutant melanoma cell lines.

## Discussion

The DIA-MS measurements in our study enhanced the detection of UPR associated proteins, many of which are low abundant. While previous LC-MS/MS targeted methods could detect and quantify transcription factors and membrane proteins critical to the UPR, these methods typically required significant development time and are often not generalizable across different instruments, making them difficult to implement widely across research groups with varying levels of expertise in proteomics^56^. To address the challenge of limited detection of UPR receptors and transducers by mass spectrometry, we instead turned to generating an expanded list of branch-specific UPR effector proteins. This approach circumvents detection limitations in mass spectrometry, enabling effective monitoring of UPR activation within global proteomics datasets.

Numerous DIA acquisition schemes have been developed to aid peptide detection^70,71^. Compared to wide-window DIA, narrow-window DIA has been shown to achieve higher peptide detection due to its enhanced specificity in precursor selection, which limits interference signals from overlapping precursors and fragments, thereby improving data interpretation. The DIA methods compared in this work had the same effective isolation window size (10 *m/z*). However, the staggered method with 20 *m/z* windows only achieved this effective isolation window using post-acquisition spectral deconvolution (**Fig. 2C**). Our DIA method comparison revealed that the semi-staggered method with a true isolation widow size of 10 *m/z* and 1 *m/z* window overlap resulted in greater peptide and protein detection, likely due to reduced MS^2^ interferences (**Fig. 2D**). Additionally, DIA was shown to result in greater log_2_ fold changes of many UPR regulated proteins compared to TMT-DDA (**Fig. 3H/I**). These differences are likely the result of the measurement of TMT reporter ions using MS^2^ spectra instead of MS^3^, potentially causing co-isolation interference and subsequent ratio compression. DIA has been previously shown to have improved quantitative accuracy over TMT-DDA^55^. The lack of selection bias during fragmentation of peptides with DIA offers greater reproducibility, higher sensitivity, and ultimately the detection of low-abundant peptides making it ideal for use in the measurement of UPR regulated proteins in different disease states, treatments, and cell lines (**Fig. 3F**).

To identify candidate branch-specific UPR targets, engineered stable cell lines (HEK293^DAX^ and Fv2e-PERK) in which the three branches of the UPR can be activated independently by small-molecules were utilized. To validate UPR regulation of these candidates, they were compared to proteins significantly upregulated in HEK293 cells treated with global UPR activators (Tg and Tm) measured via DIA-MS, in addition to an RNAseq (Tg treated) dataset resulting in a refined list of UPR branch-specific markers. Interestingly, 33 of 59 transcriptional branch-specific UPR targets outlined by Grandjean and colleagues (2019) were not identified in our proteomics analysis or were not determined to be branch-specific targets, highlighting the unique translational regulation of the UPR. Our multi-tier validation approach identified 34 new branch-specific proteomics UPR targets which were not identified via RNAseq, revealing proteomics analysis can provide important insights inaccessible by transcriptomics. Additionally, 25 genes were identified by both transcriptomics and proteomics as branch specific.

The analysis of our proteomics datasets also allowed for new assignment of genes to specific UPR branches. While the regulation of the hexosamine biosynthetic pathway (HBP) by the UPR has been previously described, we provide additional evidence of its regulation by XBP1s specifically. The HBP generates UDP-GlcNAc, a metabolite essential for N- or O-linked glycosylation. In this study, we observe significant upregulation of key enzymes—GFPT1, PGM3, and UAP1—among which UAP1 is cooperatively regulated, playing a crucial role in HBP. Furthermore, proteins from the COPI and COPII families were identified to be regulated by the IRE1/XBP1s branch. COPII coats vesicles involved in anterograde trafficking of newly synthesized secretory proteins from the ER to the Golgi, whereas COPI coated vesicles mediate the retrograde transport of escaped ER residents from the Golgi to the ER and intra-Golgi transport. Maintenance of ER-to-Golgi COPII trafficking is essential for cellular homeostasis, as previously demonstrated^72^. Additionally, COPII vesicle trafficking has been shown to be coupled to nutrient availability through the XBP1s branch of the UPR^73^. We also observed upregulation of COPII trafficking machinery in the presence of two ER global stressors (Tg and Tm). Additionally, numerous TMED proteins (TMED2, TMED5, TMED9), known to interact with coatomer (COP) complex proteins to facilitate cargo selection and vesicle formation during anterograde transport^61–63^ were found to be branch-specifically regulated by XBP1s. However, TMED proteins remain understudied, specifically in the context of the UPR^74,75^.

Several proteins significantly enriched in the activated engineered stable cell lines measured by DIA-MS were not recapitulated with either Tg or Tm treatment (**Fig. 4A**). Activation of all three branches of the UPR by Tg and Tm likely leads to crosstalk between the branches, resulting in expression patterns that deviate from those observed in engineered stable cell lines. In particular, PERK has been previously shown to impact the transcriptional and translational response of a subset of ATF6 and XBP1s target genes^76–78^. Thus, the engineered stable cell lines utilized in this study provide a unique opportunity to interrogate the transcriptional and translational targets of each branch of the response, independent of global stress and intra-branch regulation. Alternatively, it is possible that overexpression of UPR transcription factors can result in off-target DNA binding. To counteract this possibility, we applied stringent filtering using multiple datasets (transcriptomics and proteomics) to determine if a DIA-MS detected protein is UPR regulated.

Several PERK targets identified in this study (e.g. SLC1A4, EIF4EBP1, ASNS, and SESN2) were previously utilized to measure activation of the Integrated Stress Response (ISR) in both RNA-seq and proteomics (TMT-MS) datasets in Vanishing White Matter (VWM) disease^41^. VWM disease, resulting from loss-of-function mutations in eIF2B, leads to a chronically stressed phenotype akin to what is observed when eIF2α is phosphorylated by PERK^79^. Additionally, proteins involved in cellular amino acid biosynthetic processes (e.g. ASNS, CTH, PSAT1, and PSPH) and amino acid transport (e.g. SLC3A2) which were identified as PERK targets in our study were found to be significantly upregulated in a mouse model of VWM harboring the Eif2b5^R191H/R191H^ mutation. These findings confirm these proteins as targets of PERK/ISR in disease models.

Additionally, our proteomics approach confirmed numerous aminoacyl-tRNA synthetases as branch-specific targets of PERK (e.g. ASNS, CARS1, WARS1, etc.). Broadly, these enzymes are involved in the pairing of tRNAs with their cognate amino acids^80^. In the context of the UPR, ATF4 upregulates aminoacyl-tRNA synthetases, resulting in heightened protein synthesis, oxidative stress, and reduced cell viability, irrespective of CHOP’s presence^81^. Knockdown of ribosomal genes (RPL24 and RPL7) can prevent ATF4 and CHOP from increasing protein synthesis resulting in increased cell survival^82^. More recently, however, Liu and colleagues proposed that CHOP expression in cells with low level ER stress restores protein synthesis to maximize activation of all three branches of the UPR to promote cell recovery and adaptation over cell death^81^. Thus, the upregulation of aminoacyl-tRNA synthetases may reflect either a pro-adaptive or pro-death response, highlighting the complexity of the PERK branch of the UPR. Overall, our dataset reveals a comprehensive list of PERK specific markers which can be utilized to determine the state of the response in global proteomics measurements.

Application of these target proteins to evaluate the state of the UPR in four different BRAF-mutant melanoma cell lines treated with BRAF inhibitors subsequently revealed differential use of the UPR. Yang and colleagues previously demonstrated in BRAF-mutant melanoma through single-cell transcriptomics that PERK stress signaling serves as a non-genetic mechanism by which melanoma cells treated with dabrafenib, an alternative BRAF inhibitor to vemurafenib used in this study^83^, escape apoptosis within the first 72 hours of treatment. In contrast, we observed XBP1s activation using steady-state proteomics in A375 and SKMEL28 cell lines after 8 days of PLX4720 (**Fig. 7C and Supplemental Fig. 9B**). Transcriptomic analysis additionally revealed upregulation of XBP1s activity after 8 days of PLX4720 treatment, which was consistent with the findings from our proteomics dataset (**Fig. 7E**). However, while PERK activity was upregulated at the transcriptomic level, this was not reflected at the proteomic level. These findings thus emphasize that measuring UPR activity with transcriptomics alone can lead to misleading conclusions about UPR activation, as relying on transcript levels of UPR target genes may cause false assumptions about the functional importance of a gene in the translational response.

One potential mechanism by which the UPR is modulated in BRAF-mutant melanoma is through activated calcium signaling. Specifically, as reported earlier, A375 cells treated with BRAFi for 3 days showed a significant reduction in free ER Ca^2+^ content^34^. Proper calcium concentration in the ER is critical to the activity of protein folding machinery^84,85^. As discussed, the abundance of XBP1s and ATF6 target proteins significantly increased after 3 days and 3 and 8 days of BRAFi treatment, respectively, in the A375 cell line (**Fig. 7C**). This suggests that ER calcium reductions may be a critical driver of UPR activation. In general, BRAFi induced activation of IRE1/XBP1s corresponded with increased Ca^2+^ signaling activity as reported previously^34^. The cell lines reported here with XBP1s activation (A375/SKMEL28) also have much greater Ca^2+^ signaling activity in the drug tolerant state compared to those without (SKMEL5/WM88). An interesting possibility is that robust Ca^2+^ signaling activity in these cell lines may increase the efflux rate of Ca^2+^ from the ER, favoring reduced free Ca^2+^ content and induction of IRE1/XBP1s activation. This may be further supported and amplified by the stark reduction in the ER refilling mechanism, store-operated calcium entry (SOCE), also shown previously^34^. If restoration of SOCE activity in cell lines with XBP1s activation can reinstate sensitivity to BRAFi, this finding would directly link the UPR to differential drug sensitivities via a previously unconsidered Ca^2+^ mechanism, highlighting the UPR as a potential targetable pathway. Overall, our data suggests activation of the IRE1/XBP1s branch of the UPR serves as a non-genetic adaptation to promote BRAF-mutant melanoma survival on a cell-specific basis.

In conclusion, the UPR is a challenging pathway to study as different cell-types and diseases have been shown to uniquely modulate the response even when activated by an identical ER-stress-inducing treatment^36^. Additionally, the transcriptional responses initiated by the three arms of the UPR have been shown to differ between cell-types^35,86,87^. The proteomics targets we evaluated in HEK293 cells were applied in the analysis of UPR in alternative cell variants such as BRAF-mutant melanoma cell lines from different genetic backgrounds. Despite the UPR’s high clinical relevance, the specific components of its branches at the proteomics level have been largely undefined until now. Our study provides a valuable resource for investigating the UPR branch-specifically using proteomics (**Supplemental Table S14)**.

## Acknowledgments

We would like to thank Dr. Lee Cantrell for his advice on DIA-MS and data analysis. Dr. Minsoo Kim (Vanderbilt University) for his thoughtful edits and figure suggestions. Additionally, we would like to thank Dr. Luke Wiseman (Scripps Research Institute) for providing us with the HEK293^DAX^ and HEK293 Fv2e-PERK engineered stable cell lines.

## Data Availability

The R scripts used in the analysis of TMT-DDA and DIA data are available on GitHub (https://github.com/Plate-Lab/). The mass spectrometry proteomics data are deposited to the ProteomeXchange Consortium via the PRIDE partner repository under the accession code PXD061073. All other necessary data are contained within the manuscript or can be shared by the Lead Contact upon request.

## Supplemental data

This article contains supplemental data.

S1. List of branch-specific RNAseq UPR targets with accession numbers from Grandjean et al., 2019.

S2. Unnormalized TMT-DDA data, filtered at two peptides per protein.

S3. Log_2_ transformed and median normalized HEK293^DAX^ data measured by DIA.

S4. Log_2_ transformed and median normalized Fv2e-PERK data measured by DIA.

S5. Measurement of ATF6 RNAseq targets by DIA and TMT-DDA MS.

S6. Identification of proteomics branch-specific ATF6 targets.

S7. Measurement of XBP1s RNAseq targets by DIA and TMT-DDA MS.

S8. Identification of proteomics branch-specific XBP1s targets.

S9. Log_2_ transformed and median normalized HEK293 cells with global UPR activation measured by DIA.

S10. Proteins determined to be cooperatively regulated or not branch-specific.

S11. Measurement of PERK RNAseq targets by DIA and TMT-DDA MS.

S12. Identification of proteomics branch-specific PERK targets.

S13. Log_2_ transformed and median normalized DIA data collected from four BRAF-mutant melanoma cell lines treated for 0, 3, and 8 days with BRAF-inhibitor.

S14. List of branch-specific proteomics UPR targets (accession number and gene symbol).

