## Supplemental Information for "Dissecting Branch-Specific Unfolded Protein Response Activation in Drug-Tolerant BRAF-Mutant Melanoma using Data-Independent Acquisition Mass Spectrometry"

For

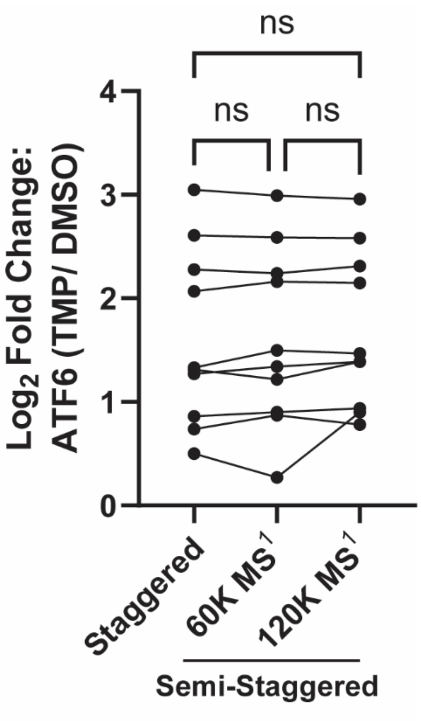


**Supplemental Figure 1.** ***Quantification of ATF6 proteomics targets using three different DIA methods***. Staggered (20 *m/z* isolation window with 10 *m/z* overlap) and two semi-staggered methods (10 *m/z* isolation window with 1 *m/z* overlap) with precursor scans taken at two $MS^1$ resolutions (60,000 and 120,000) were compared. Quantification of proteins was not significantly different between DIA methods. Statistical analysis was performed using one-way ANOVA test with Tukey's post hoc test. A $p$ value < 0.05 was considered statistically significant.

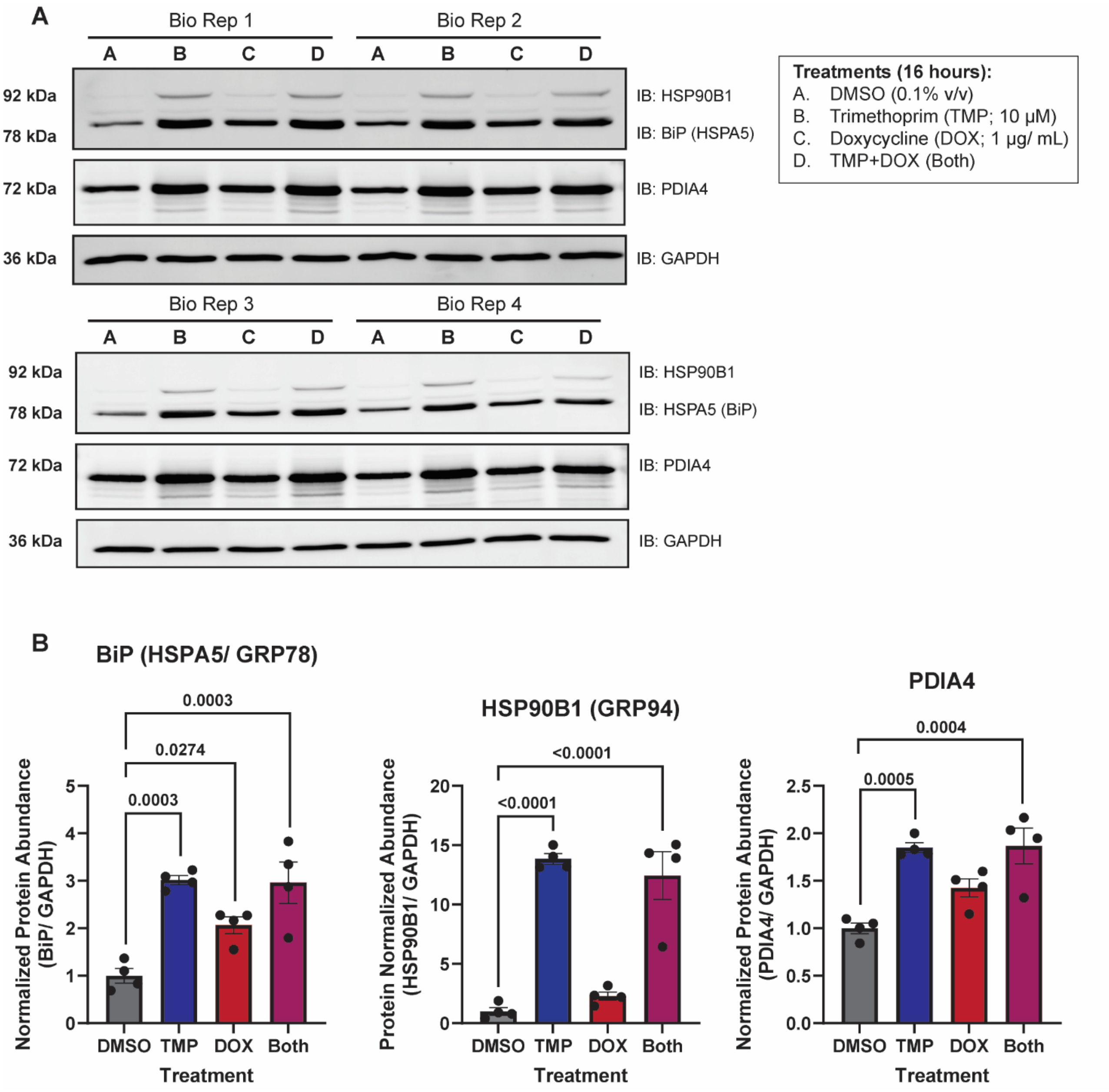


**Supplemental Figure 2.** ***Western blot validation of ATF6 activation in HEK293$^{DAX}$ cells.*** (**A**) Western blots of ATF6 protein markers (HSP90B1 (GRP94), BiP (HSPA5/GRP78), and PDIA4) in HEK293$^{Dax}$ cells treated with DMSO (0.1%), TMP (10 µM), DOX (1 µg/mL), or both (TMP (10 µM) and DOX (1 µg/mL)) for 16 hours used for the analysis ATF6/XBP1s pathways via TMT-DDA and DIA LC-MS/MS analysis. GAPDH was used as a housekeeping gene for loading control. Displayed blot sections are from the same blot image and exposure settings. (**B**) Quantification of western blots in panel (A; BiP, HSP90B1, and PDIA4), normalized to GAPDH band intensities and compared to the DMSO control (vehicle). n =4, mean ± SEM. Statistical analysis was performed using a one-way ANOVA test with Tukey's post hoc test was computed. A *p* value < 0.05 was considered statistically significant.

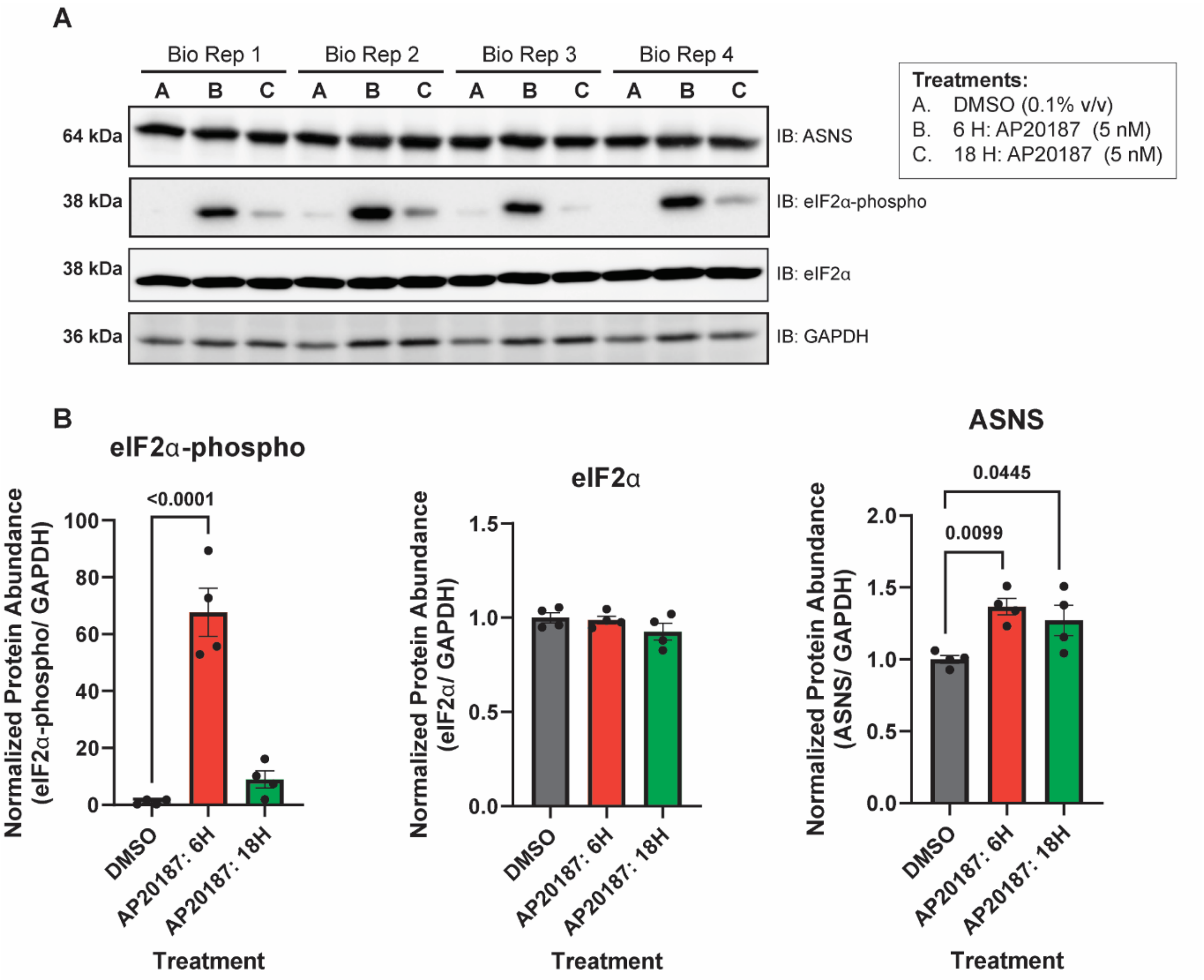


**Supplemental Figure 3.** ***Western blot validation of PERK activation in Fv2e-PERK cells.*** (**A**) Western blots of PERK protein markers (eIF2α-phospho, eIF2α, and ASNS (asparagine synthase) in Fv2e-PERK cells treated with DMSO (0.1%) for 18 hours and AP20187 (5 nM) for either 6 or 18 hours used in the analysis of the PERK pathway via TMT-DDA and DIA LC-MS/MS analysis. GAPDH was used as a housekeeping gene for loading control. Displayed blot sections are from the same blot image and exposure settings. (**B**) Quantification of western blots in panel (A; eIF2α-phospho, eIF2α, and ASNS), normalized to GAPDH band intensities and compared to the DMSO control (vehicle). n =4, mean ± SEM. Statistical analysis was performed using a one-way ANOVA test with Tukey's post hoc test was computed. A $p$ value < 0.05 was considered statistically significant.

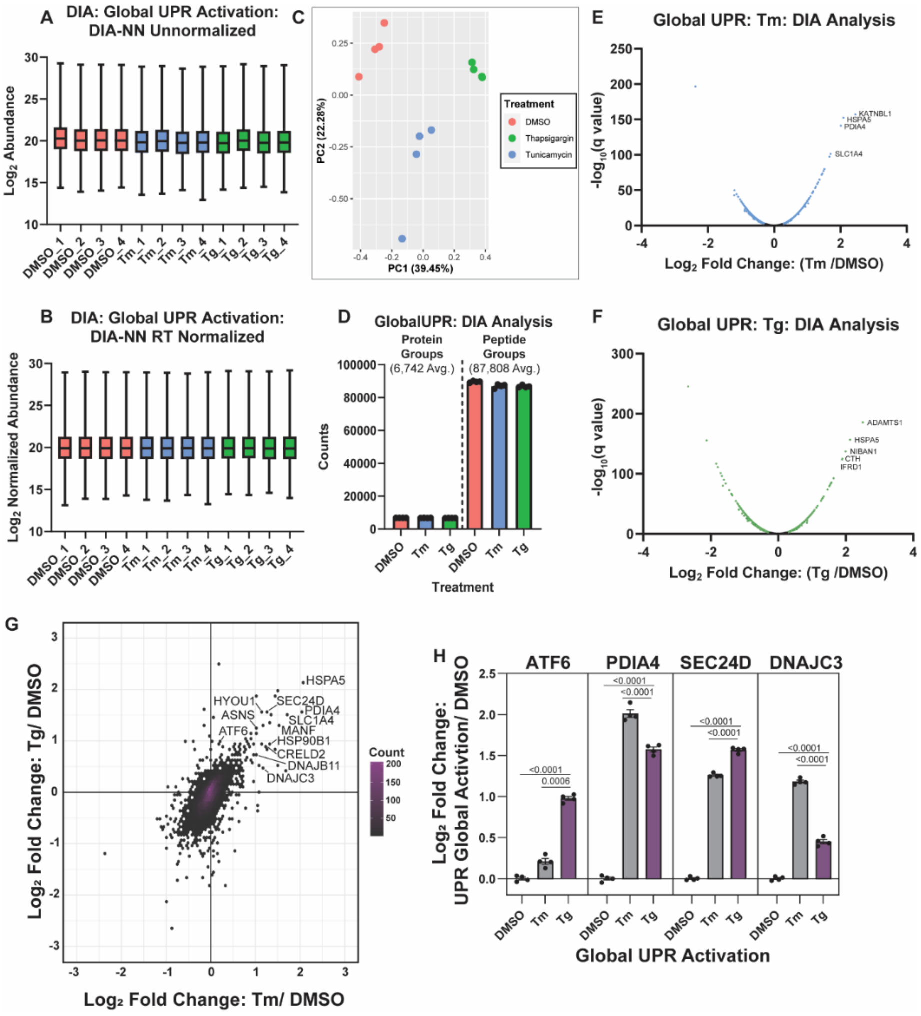


**Supplemental Figure 4.** ***Validation of the HEK293 global UPR activation DIA-MS dataset.*** Box and whisker plots showing the distribution of protein abundances pre (**A**) and post (**B**) retention time (RT) median normalization, implemented in DIA-NN of HEK293 cells treated with either DMSO (0.1%), Tm (500 nM), or Tg (500 nM) for 16 hours. Box and whisker plots show median, 25$^{th}$ and 75$^{th}$ quartiles, and minimum and maximum values. (**C**) Principal component analysis (PCA) plot generated to ensure biological replicates (n= 4) cluster by treatment. (**D**) Bar plot showing the number of protein groups and peptide groups detected in the globally activated UPR samples. (**E-F**) Volcano plots of HEK293 cells activated with Tm (500 nM) (**E**) and Tg (500 nM) (**F**), measured by DIA-MS. Colored points represent proteins with a $-\log_{10}$(q value) $\geq 2$. **(G)** Correlation analysis comparing the $\log_2$ fold change of HEK293 cells treated with Thapsigargin (Tg; 500 nM) or Tunicamycin (Tm; 500 nM) for 16 hours compared to DMSO. The black diagonal dashed line indicates y=x. **(H)** Bar plots, of UPR regulated proteins, showing the $\log_2$ fold changes of HEK293 cells treated with Tg and Tm compared to DMSO. n=4, mean ± SEM. Statistical analysis was performed using one-way ANOVA per protein and post hoc Tukey's multiple comparison test was used to test significance, $p < 0.05$ considered statistically significant.

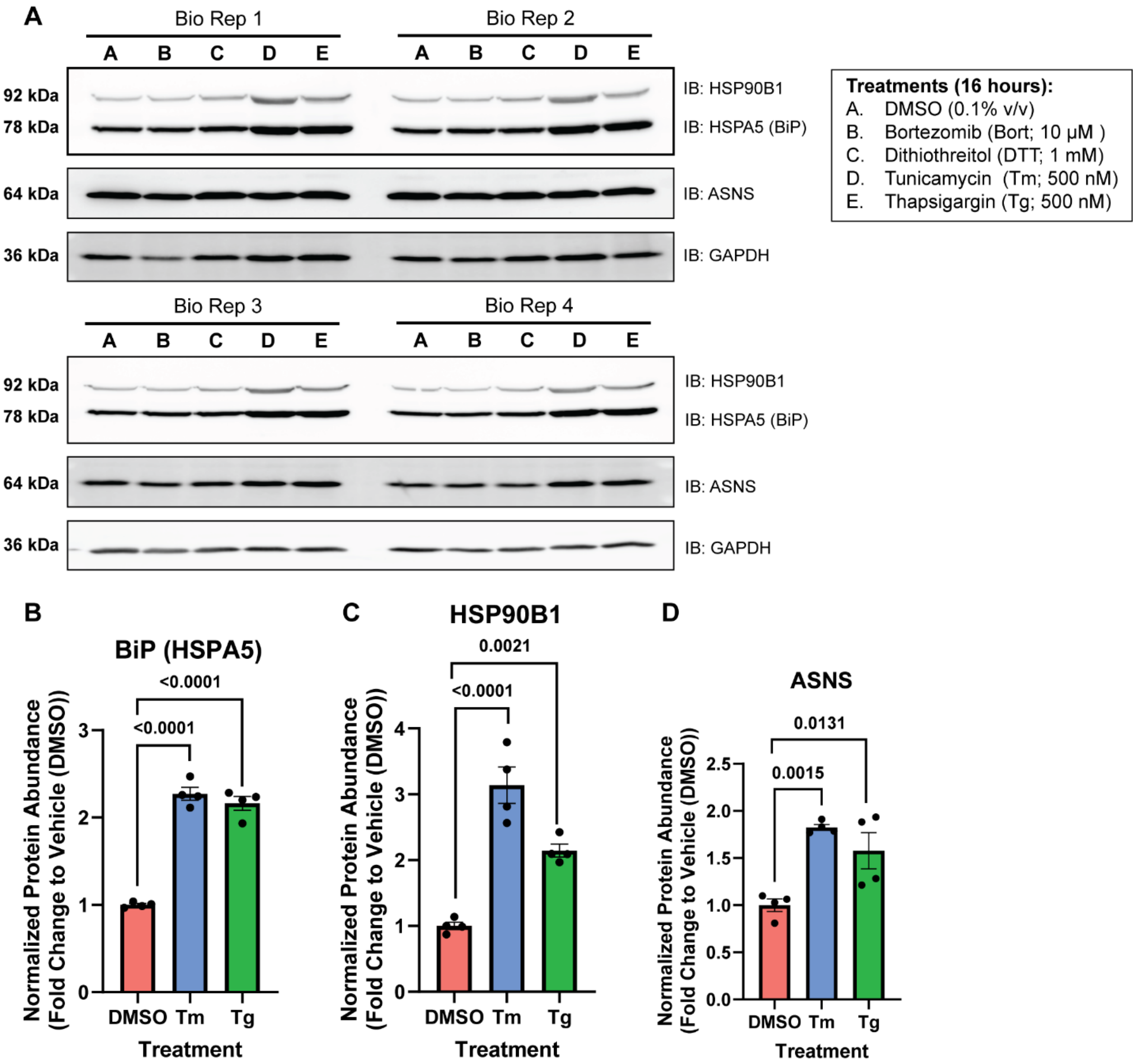


**Supplemental Figure 5.** ***Western blot validation of HEK293 global UPR activation.*** (**A**) Western blots of UPR protein markers (HSP90B1 (GRP94), BiP (HSPA5/GRP78), and ASNS) in HEK293 cells treated with DMSO (0.1%), bortezomib (10 µM), dithiothreitol (1 mM), Tm (500 nM) and Tg (500 nM) for 16 hours used in the analysis global UPR activation via DIA LC-MS/MS analysis. GAPDH was used as a housekeeping gene for loading control. Displayed blot sections are from the same blot image and exposure settings. (**B-D**) Quantification of western blots in panel (a), normalized to GAPDH band intensities and compared to the DMSO control (vehicle). n=4, mean ± SEM. Statistical analysis was performed using a one-way ANOVA with Benjamini, Krieger, and Yekutieli multiple testing correction, p <0.05 considered statistically significant.

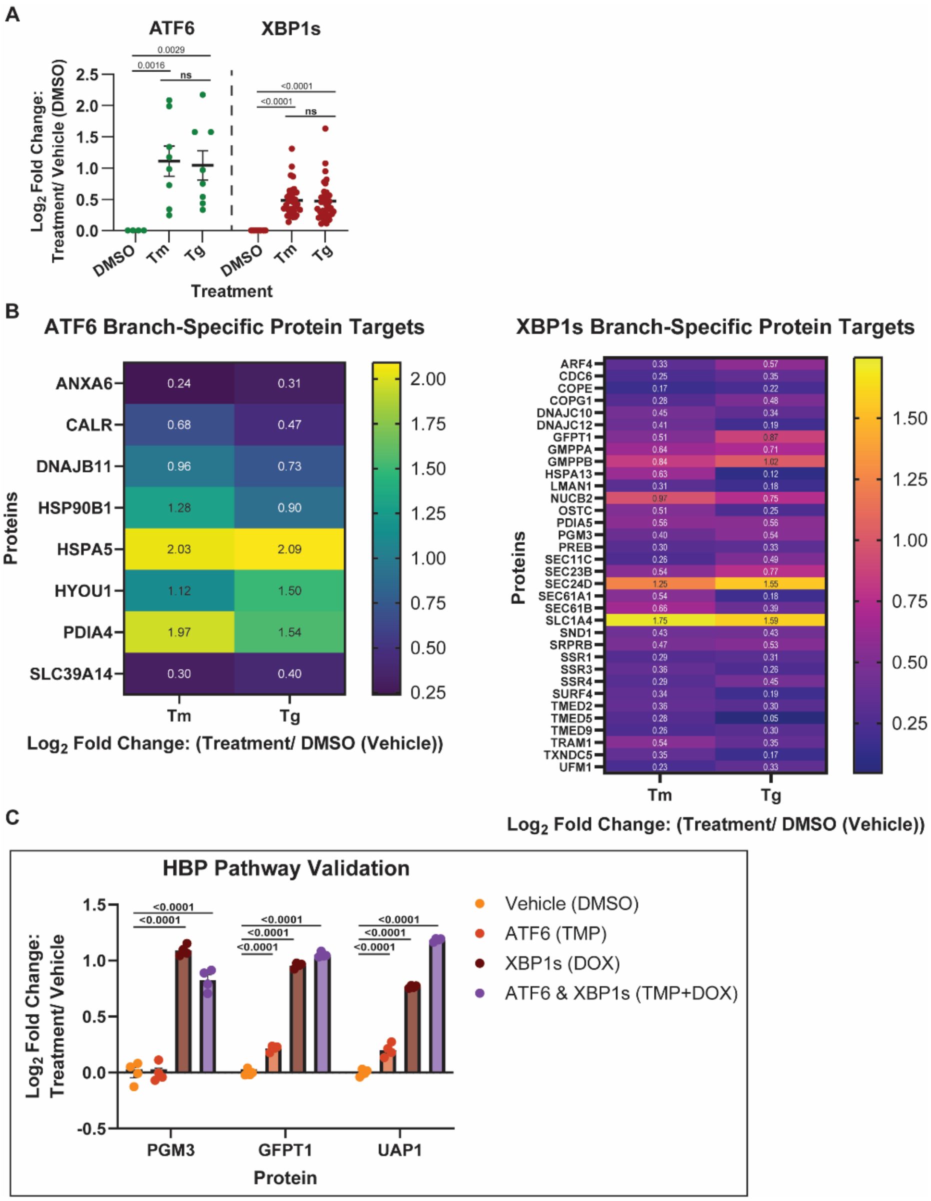


**Supplemental Figure 6.** ***Quantification of branch-specific ATF6 and XBP1s targets in HEK293 cells treated with global UPR activators.*** (**A**) Proteomics UPR targets shown in (A), for ATF6 and XBP1s, measured in HEK293 cells activated with either Tm or Tg. n =4, mean ± SEM. Statistical analysis was performed using one-way ANOVA with Benjamini, Krieger, and Yekutieli multiple testing correction, p <0.05 considered statistically significant. (**B**) Heatmap of the quantification of branch-specific ATF6 and XBP1s proteins in Tm and Tg global UPR activated samples. Values within cells represent the $log_2$ fold change of the treatment by DMSO (vehicle) control. (**C**) Quantification of the enzymes involved in the hexosamine biosynthesis pathway (HBP) in ATF6, XBP1s, and TMP+DOX treated $HEK293^{DAX}$ cells. n =4, mean ± SEM. Statistical analysis was performed per protein using one-way ANOVA with Benjamini, Krieger, and Yekutieli multiple testing correction, p <0.05 considered statistically significant.

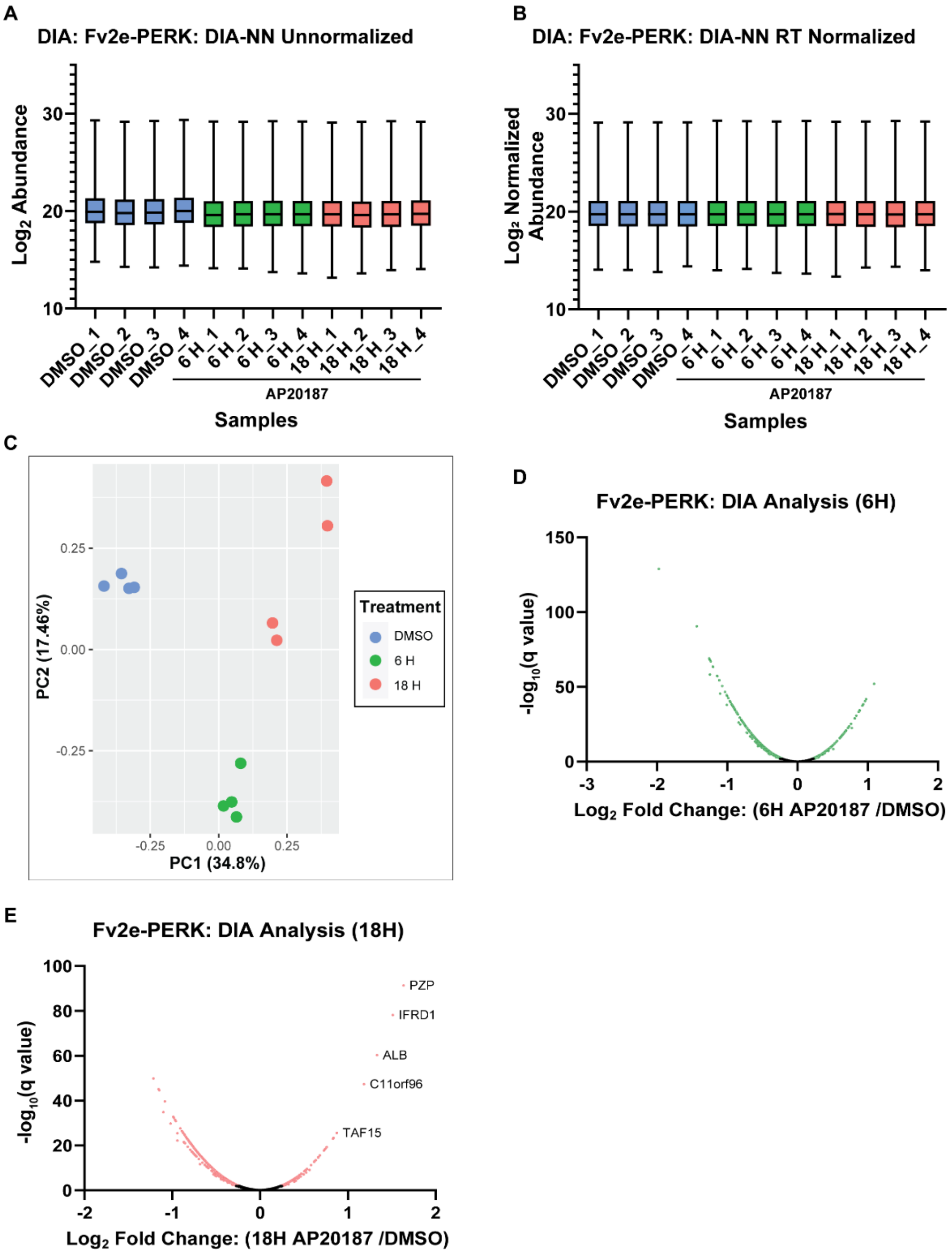


**Supplemental Figure 7.** ***Validation of the Fv2e-PERK DIA-MS dataset.*** (**A-B**) Box and whisker plots showing the distribution of protein abundances pre (a) and post (b) retention time (RT) median normalization, implemented in DIA-NN of Fv2e-PERK cells treated with DMSO (0.1%) for 18 hours and AP20187 (5 nM) for either 6 or 18 hours. Box and whisker plot shows median, 25$^{th}$ and 75$^{th}$ quartiles, and minimum and maximum values. (c) Principal component analysis (PCA) plot generated to ensure biological replicates (n= 4) cluster by treatment. (**D-E**) Volcano plots of Fv2e-PERK cells activated with AP201878 for 6 hours (d) and 12 hours (e), measured by DIA-MS. Colored points represent proteins with a -log10(q value) $\geq$ 2.

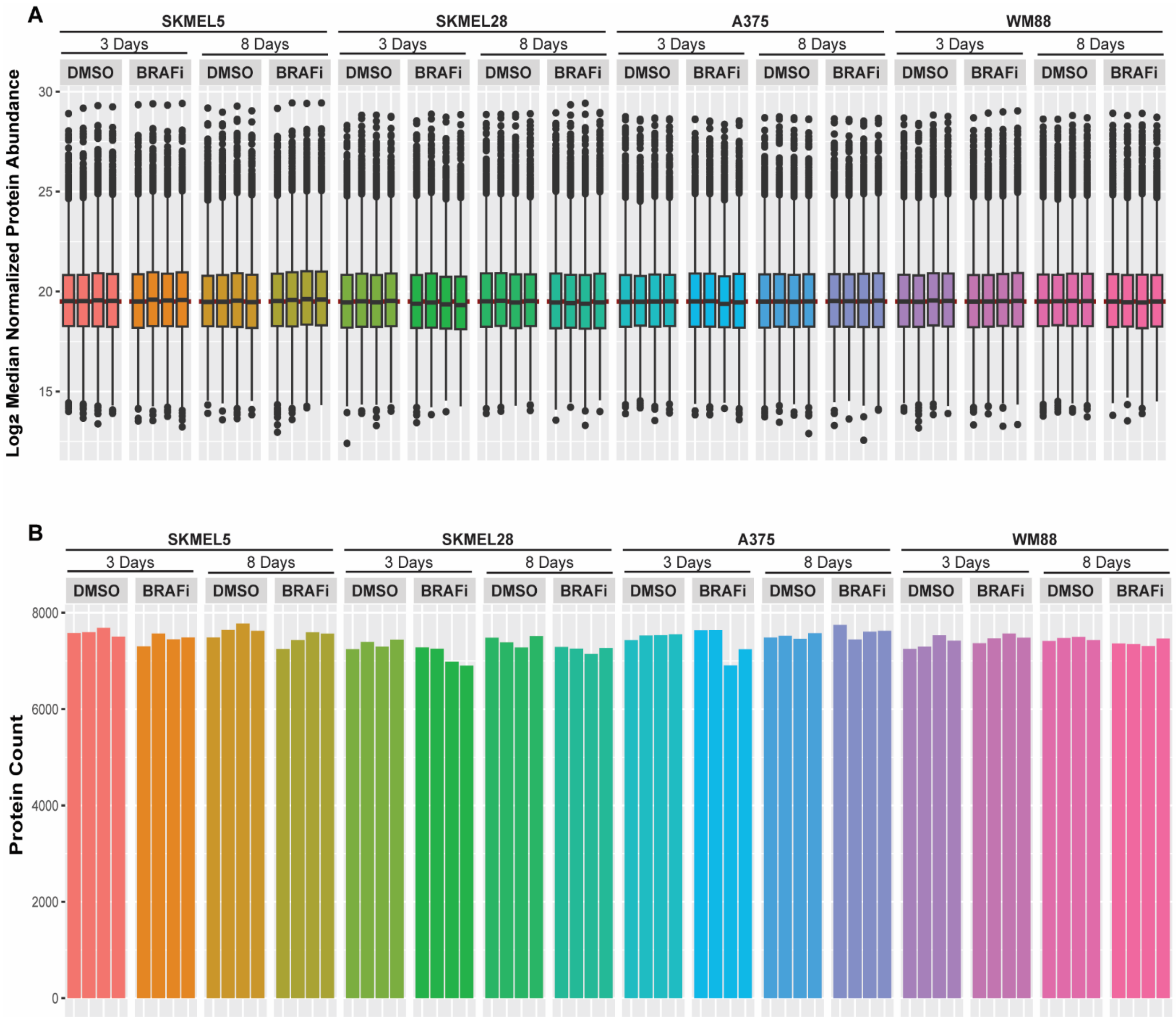


**Supplemental Figure 8.** ***Validation of the LC-MS/MS proteomics data collected on BRAFi treated (3 and 8 days) BRAF-mutant melanoma cell lines (SKMEL5, SKMEL28, A375, and WM88).*** (**A**) Distribution of protein abundances per sample grouped by cell line and treatment. (**B**) Number of proteins (2 peptides per protein) identified per sample grouped by cell line and treatment.

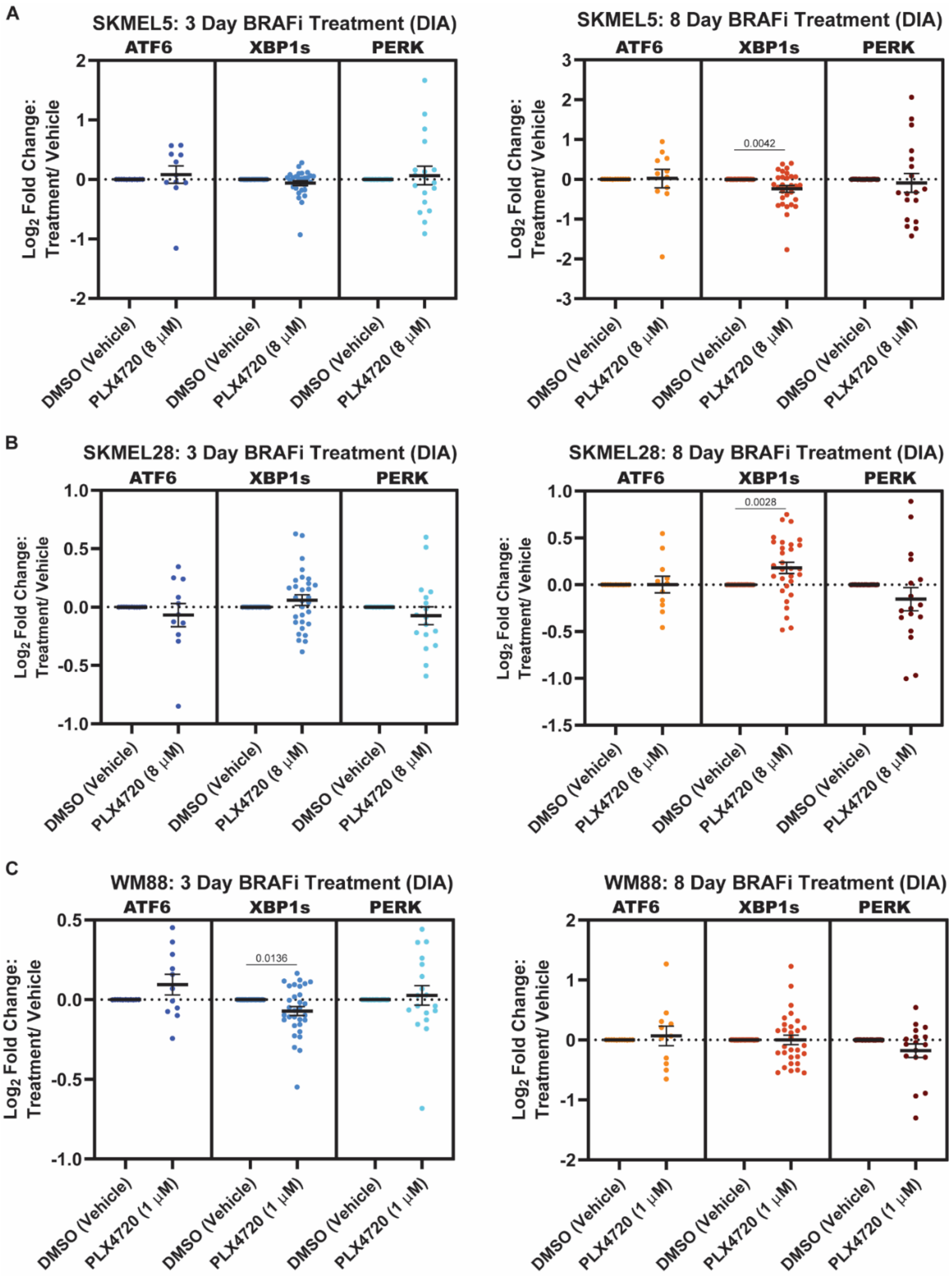


**Supplemental Figure 9.** ***UPR status in BRAFi treated BRAF-mutant melanoma.*** Evaluation of branch-specific UPR activation in **(A)** SKMEL5, **(B)** SKMEL28, and **(C)**WM88 BRAF-mutant melanoma treated for 3 and 8 days with BRAF inhibitor using LC-MS/MS (DIA) proteomics targets.
